# Adaptive therapy for ovarian cancer: An integrated approach to PARP inhibitor scheduling

**DOI:** 10.1101/2023.03.22.533721

**Authors:** Maximilian Strobl, Alexandra L. Martin, Jeffrey West, Jill Gallaher, Mark Robertson-Tessi, Robert Gatenby, Robert Wenham, Philip Maini, Mehdi Damaghi, Alexander Anderson

## Abstract

Toxicity and emerging drug resistance are important challenges in PARP inhibitor (PARPi) treatment of ovarian cancer. Recent research has shown that evolutionary-inspired treatment algorithms which adapt treatment to the tumor’s treatment response (adaptive therapy) can help to mitigate both. Here, we present a first step in developing an adaptive therapy protocol for PARPi treatment by combining mathematical modelling and wet-lab experiments to characterize the cell population dynamics under different PARPi schedules. Using data from *in vitro* Incucyte Zoom time-lapse microscopy experiments and a step-wise model selection process we derive a calibrated and validated ordinary differential equation model, which we then use to test different plausible adaptive treatment schedules. Our model can accurately predict the *in vitro* treatment dynamics, even to new schedules, and suggests that treatment modifications need to be carefully timed, or one risks losing control over tumour growth, even in the absence of any resistance. This is because our model predicts that multiple rounds of cell division are required for cells to acquire sufficient DNA damage to induce apoptosis. As a result, adaptive therapy algorithms that modulate treatment but never completely withdraw it are predicted to perform better in this setting than strategies based on treatment interruptions. Pilot experiments *in vivo* confirm this conclusion. Overall, this study contributes to a better understanding of the impact of scheduling on treatment outcome for PARPis and showcases some of the challenges involved in developing adaptive therapies for new treatment settings.

## 1. Introduction

PARP (poly-adenosine ribose polymerase) inhibitors (PARPis) are revolutionizing the landscape of ovarian cancer treatment. These small molecule inhibitors target the PARP family of proteins, in particular PARP1 and PARP2, which help to detect single strand DNA (ssDNA) damage and orchestrate the subsequent repair^3,4^. This results in a buildup of ssDNA breaks which interfere with DNA regulation and replication (Figure 1a). If the cell attempts to divide, then the replication fork will stall at ssDNA breaks, which causes cell cycle arrest, and double strand breaks^2,4,5^ (DSBs). Furthermore, there is growing evidence that PARPis additionally promote this process by trapping PARP proteins directly on the DNA, leaving further obstacles for the cell to resolve^4^ (Figure 1a). While the damage caused by PARPis can be repaired via the homologous repair (HR) pathway, HR is deficient in many ovarian tumors due to, for example, mutations in BRCA1 or BRCA2. As a result, ovarian cancers rely on more error prone backup mechanisms, such as non-homologous end joining, making PARPis an effective treatment option^2,4-7^ (an effect referred to as “synthetic lethality”).

**Figure 1.**
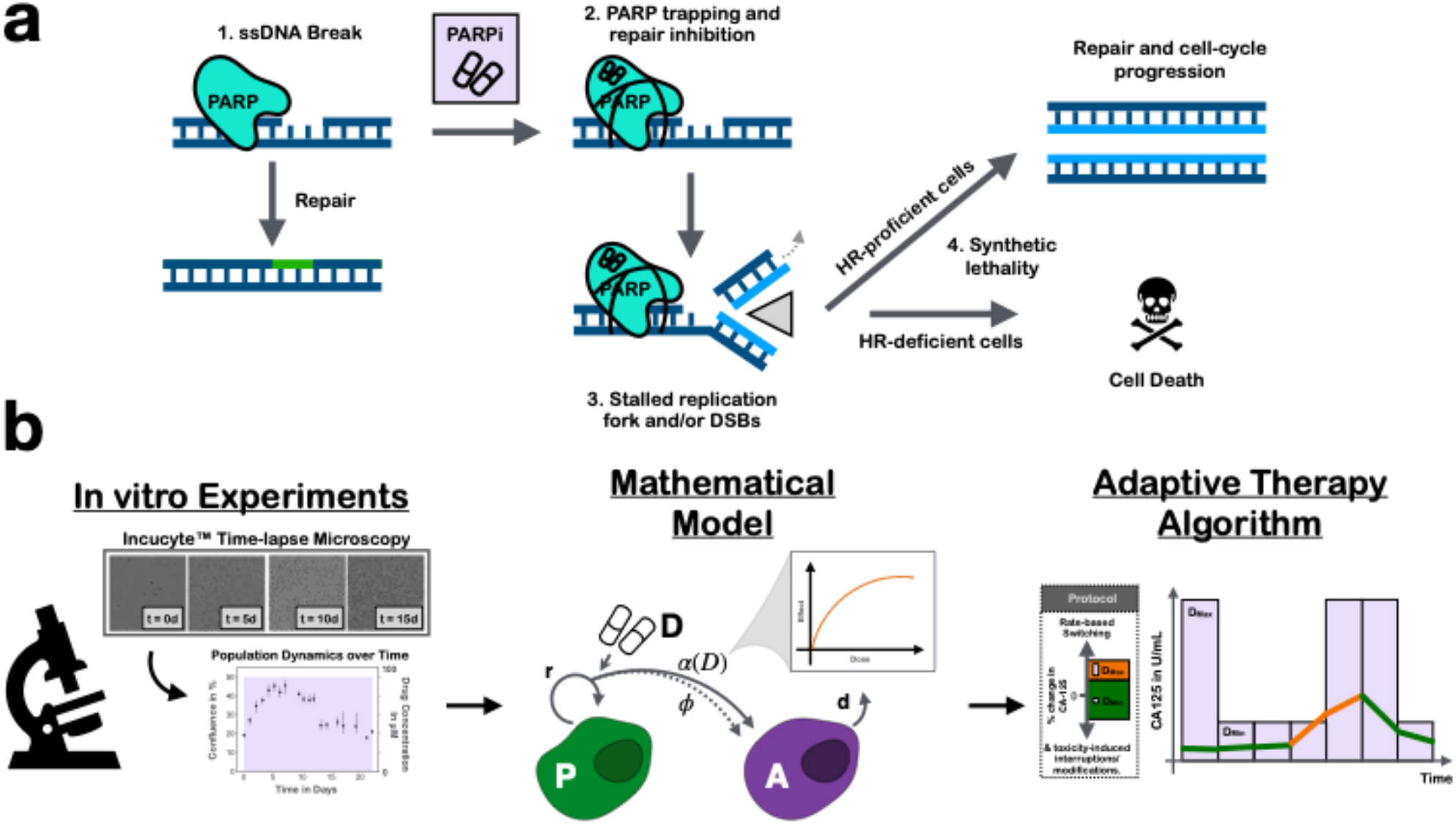
PARPis are revolutionizing ovarian cancer treatment but toxicity and developing resistance are important challenges in the clinic. Here, we developed a mathematical model to help address these issues through more personalized treatment scheduling. **a)** The mechanism of PARPi-mediated cytotoxicity. PARPs are important proteins in the repair of single strand DNA breaks. But PARPis can trap PARPs on the DNA, which results in stalled replication forks and DSBs during DNA replication. Only cells with intact HR pathways can repair this damage, but tumor cells are typically HR-deficient, and are thus killed. Redrawn with permission from Noordermeer and van Attikum ^2^. **b)** Outline of our paper. Using in vitro experiments we developed, calibrated, and validated a mathematical model of PARPi treatment. Subsequently we used this model to explore plausible adaptive treatment algorithms.

Historically, ovarian cancer has been a particularly challenging disease to treat because the majority of patients (70% ^8^) are diagnosed with stage III or IV disease, and because it is remarkably adept at overcoming treatment. Up to 80% of patients recur after first-line therapy^9^ and for many years the 5-year survival rate has remained at only around 30%^10^. But, thanks to PARPis this picture is promising to change. In this study, we will focus on Olaparib (AstraZeneca) which is the longest approved and one of the most widely used of the three currently approved agents. It is given orally and is primarily used as maintenance therapy, which means that treatment is administered after chemotherapy has been completed, with the aim to eradicate the disease or, at least, to push back progression^4^. Its benefit in patients has been demonstrated in multiple clinical trials^11-15^, most prominently the SOLO1 (NCT01844986) and SOLO2 (NCT01874353) phase III studies. For example, SOLO2 evaluated single agent Olaparib maintenance therapy for up to 2 years after chemotherapy in recurrent patients with BRCA mutations, and found that compared to placebo it increased median progression free survival (PFS) by 13.6 months^14,15^ and median overall survival by 12.9 months. But, in both studies most patients still saw their tumor recurring within 5 years^14-16^. Furthermore, around 40% of patients required dose adjustments due to serious Grade 3 or 4 adverse events, such as anaemia^17^. As such, there is an important need to investigate how we can administer Olaparib more safely and effectively.

An approach which is establishing itself as a powerful way to improve treatment scheduling is mathematical modeling^18-21^. A key challenge in optimizing treatment protocols is the large number of plausible schedules to investigate, and the fact that tumors are complex systems in which processes on different time and spatial scales interact. Mathematical models allow for the systematic interrogation of different mechanisms, and unlike traditional laboratory models, can typically provide outputs in hours rather than days or weeks, and are cheap to run. For example, Meille, et al. ^22,23^ recently used mathematical modeling to review scheduling of docetaxel and epirubicin combination treatment in metastatic breast cancer patients, after a previous trial had resulted in lethal side effects. The resulting model-optimized regimen was found to be significantly better tolerated in a subsequent Phase I/II study and showed promise of improved response^22,23^. Mathematical modeling has also been used to optimize treatment in preclinical models of ovarian cancer (see Botesteanu, et al. ^24^ for an in-depth review). For example, Jain and Meyer-Hermann ^25^ developed a model that connected carboplatin-induced DNA damage with anti-apoptotic Bcl-xL signaling in order to investigate how to optimally combine Bcl inhibitors with carboplatin chemotherapy. Calibrating their model with *in vivo* data, they predicted successfully that giving the novel Bcl inhibitor ABT-737 during, or after, chemotherapy gave the best results^25^. Similarly, others have used modeling to study the scheduling of paclitaxel chemotherapy^26^, the best order of chemotherapy and surgery in first-line treatment^27,28^, or the optimal sequencing of targeted therapies for ovarian cancer^29,30^.

In addition to improving the efficacy and tolerability of treatment, mathematical modeling has provided new perspectives on the management of drug resistance. Current treatment regimens are designed to maximize cell kill in an attempt to cure. However, in light of our growing understanding of the complex, evolving nature of a tumor, mathematical modeling has shown that a more conservative treatment may help to control resistance for longer in advanced cancers^31-37^. This is because it is thought that in these settings drug tolerant or resistant cells likely exist prior to treatment, but are suppressed by competition for space and resources with more sensitive cancer cell subpopulations^38,39^. Aggressive treatment removes this suppression and allows resistance to emerge. In contrast, a growing body of theoretical and experimental studies is showing that so-called “adaptive therapy”^32,38^, which carefully modulates treatment in a personalized fashion based on the tumor’s response dynamics so as to maintain sensitive cells and competitively suppress resistance, can help to extend PFS when cure with aggressive treatment is unlikely (e.g. refs^1,33,35-37,40,41^). A pilot Phase IIb study in androgen deprivation treatment in prostate cancer patients has demonstrated clinical feasibility of adaptive therapy^42,43^. In comparison to matched historical control patients treated with continuous, standard-of-care therapy, adaptively treated patients from this trial received 46% less treatment whilst their median time to progression was 19.2 months longer than the control cohort^43^.

These promising results raise the question of whether adaptive therapy may also help to extend PFS in PARPi maintenance therapy. Olaparib’s short half-life, oral administration, and the availability of CA-125 as an easily accessible biomarker of tumor response suggest that adaptive administration may be feasible. However, *how* should therapy be adapted? The aim of this paper is to begin answering this question by developing the first mathematical model of PARPi treatment in ovarian cancer (Figure 1b). To calibrate the model, we used *in vitro* time-lapse microscopy experiments to measure the population dynamics of ovarian cancer cells in response to PARPi treatment under different seeding conditions (low and high initial density) and treatment schedules (continuous and intermittent treatment at different drug concentrations). Leveraging these data, we systematically evaluated different plausible models of treatment response to derive our final calibrated and validated model, and to shed biological insights into the observed dynamics (Figure 2). To conclude, we used our model to explore different possible PARPi treatment algorithms, showing that strategies which adjust treatment by modulating the dose are predicted to be superior to those skipping treatments. This finding in the absence of drug resistant cells suggests that treatment modifications for PARPis need to be carefully timed and most likely should avoid dose skipping, and we conclude with a discussion of a possible adaptive PARPi strategy. Overall, our study contributes to a better understanding of the impact of scheduling for PARPis, and showcases the first steps in developing adaptive therapies in a new treatment setting.

**Figure 2.**
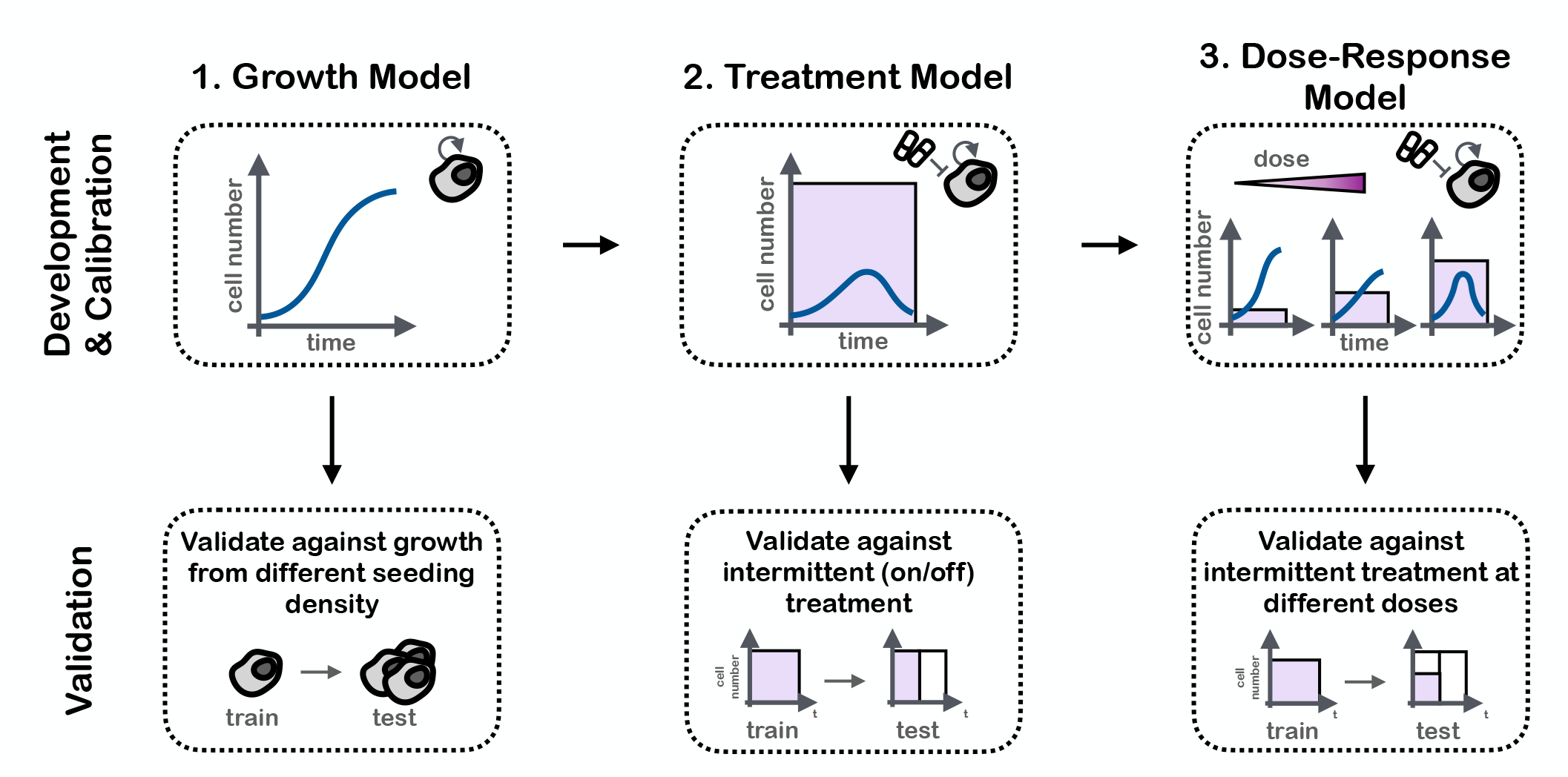
Overview of the model development process. We proceeded in a step-wise fashion, where firstly we developed a model for growth under untreated conditions. Subsequently, we extended the model to describe the time dynamics in response to continuous treatment at MTD (100µM), and finally we modeled the response to continuous treatment at varying intermediate doses. In addition, we tested the model at each step using independent data from experiments under conditions different to those under which the model was calibrated.

## 2. Methods

### 2.1. Cell Culture

OVCAR3 and OVCAR4 cells were acquired from American Type Culture Collection (ATCC, Manassas, VA, 2007 to 2010) and cultured in Roswell Park Memorial Institute (RPMI) medium (ThermoFisher) supplemented with 10% Fetal Bovine Serum and 1% penicillin/streptomycin. Every 3-4 weeks the medium was additionally supplemented with MycoZap (Lonza) to prevent mycoplasma contamination. At all times cells were kept at 37C and in a 5% CO_2_ atmosphere.

### 2.2. Measuring the drug response dynamics during continuous drug exposure

To characterize how the tumor cells grew in the absence of treatment and when exposed continuously to different drug concentrations, we seeded cells in a 48 well flat-bottom plate (Costar Corning) and left them to attach overnight in 200µL culture medium. Subsequently, we aspirated the medium and replaced it with treated growth medium, containing 0, 1, 10, 25, 50 or 100µM Olaparib (AstraZeneca), and monitored their growth for 9 days. We carried out two versions of this assay: i) a “low-density” version in which we seeded cells at 5,000 cells per well, and ii) a “high-density” version in which each well started with 60,000 cells. In each case, 3 replicates were performed for each experimental condition. During the experiment the medium was changed every 3 days. We also tested changing the medium daily but found that this did not significantly change the growth dynamics (not shown).

To prepare the treated medium, we first created a stock containing 100µM of drug by dissolving Olaparib (AstraZeneca) in 1mL Dimethyl sulfoxide (DMSO), filtering the solution using a 0.22nm syringe filter, and dissolving it in our regular culture medium. Next, we diluted this maximum tolerated dose (MTD) stock with normal culture medium to obtain batches with 1-50µM Olaparib. We verified that the DMSO did not adversely impact the cells’ growth dynamics (not shown).

### 2.3. Measuring the dynamics in response to drug withdrawal

To test how the cells responded to treatment withdrawal, we seeded 10,000 cells per well in a 48 well flat-bottom plate (Costar Corning) and left them to attach overnight in untreated culture medium. Next, we aspirated the medium and replaced it with treated medium for 0, 1, 2, 4, 7, or 21 days before we withdrew treatment again by replacing the medium with regular culture medium. We repeated this experiment twice: once where cells were treated at 50µM and once where cells were treated at 100µM Olaparib (treated medium was prepared as specified in Section 2.2.). In each case, we carried out 3 replicates for each experimental condition. During the 21 day experiment the medium was changed every 3 days, or when it was time to withdraw drug from a well.

### 2.4. Real-time imaging and data processing

Cell growth was monitored once per day using an IncuCyte ZOOM S2 time-lapse microscopy system (Essen BioScience; see Figure 3a for examples). Confluence was measured based on phase-contrast, white light images, which were analyzed using the IncuCyte ZOOM software (10x magnification; confluence estimated based on 2 images per well). On two occasions we accidentally removed large numbers of cells when aspirating out medium during medium changes, and thus we did not include measurements from these wells in our analysis. In addition, when measuring the treatment response of OVCAR4 cells under 100µM Olaparib for 21 days (protocol per Section 2.3.), we found that after 13-14 days the imaging system was greatly overestimating confluence due to the build-up of debris from dead cells on the plate. To avoid this from confounding our results, we decided not to include the data from days 15-21 in our analyses. The raw and curated data are available on our github repository, as is a Jupyter Notebook detailing every data curation/processing step (jnb_dataProcessing.ipynb).

**Figure 3.**
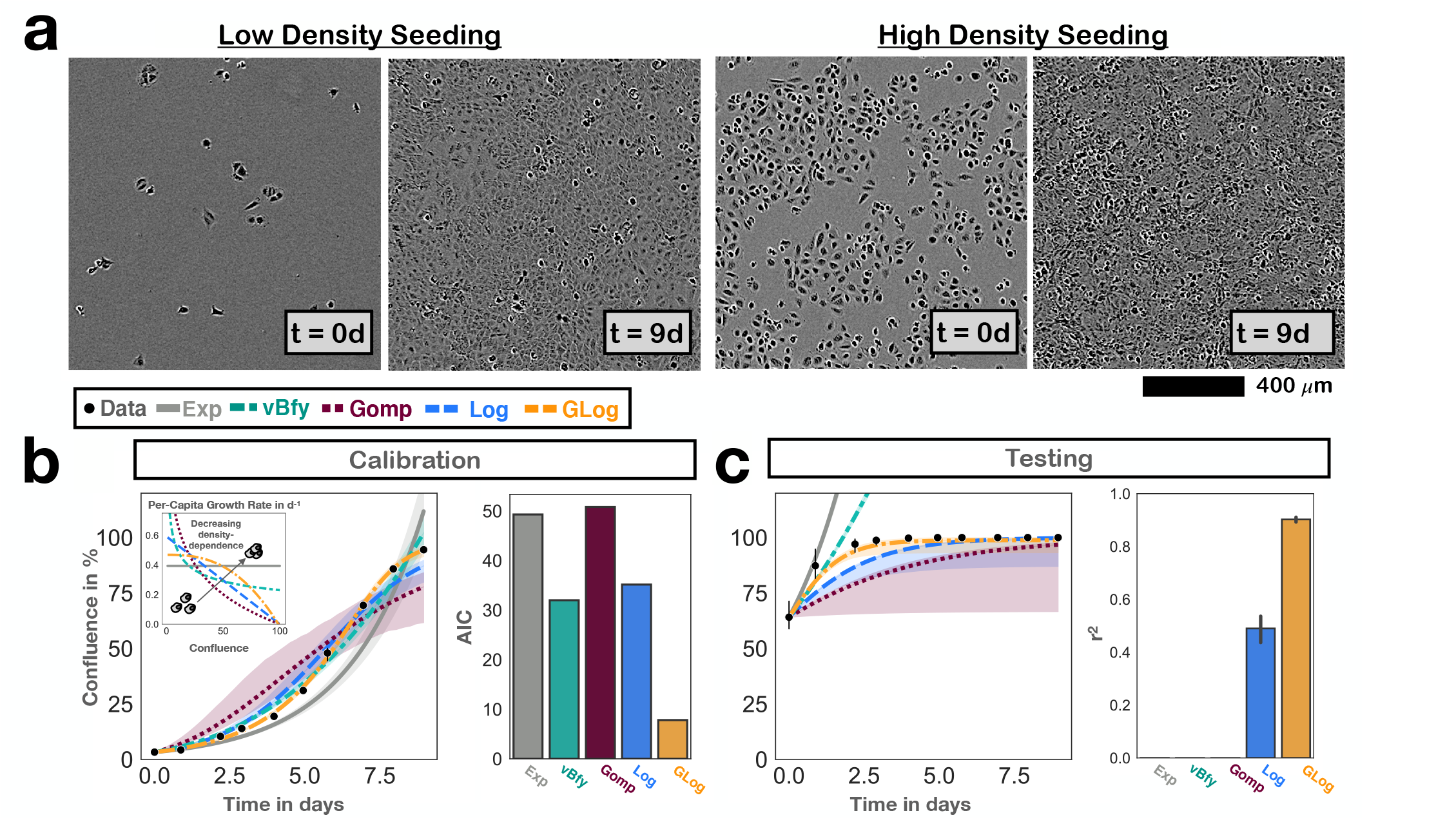
Development of the growth model to describe the population expansion in the absence of treatment. Points and bars denote mean and 95% CIs of observed confluence (n=3 independent replicates). Solid lines show the model predictions based on the maximum likelihood estimate, and bands indicate 95% CIs. **a)** Example Incucyte microscopy images based on which we assessed the growth and treatment dynamics over time. **b)** Comparison of the descriptive ability of 5 commonly used growth models in fitting the untreated growth data from cells seeded at low density (Exp: exponential; vBfy: von Bertalanffy; Gomp: Gompertz; Log: Logistic; GLog: Generalized Logistic; see Section 2.6. for equations). The GLog model achieves the best fit, even if its additional parameter is taken into account (lowest AIC score). **c)** Testing of the growth models by comparing their predictions for when cells are initially seeded at 60% confluence with the experimentally observed dynamics. This shows that the GLog model is also the most predictive model, and corroborates our choice of this growth model.

### 2.5. *In vivo* experiments

All animals were maintained in accordance with IACUC standards of care in pathogen-free rooms, in the Moffitt Cancer Center and Research Institute (Tampa, FL) Vivarium. One week before inoculation with tumor cells (5 × 10^6^ OVCAR3 cells, subcutaneously), 6–8 weeks old, female NSG mice (Charles River Laboratories) were assigned to one of the following four treatment arms: 1: Control group, treated with vehicle (DMSO) intraperitoneally. 2: MTD group, treated with PARPi (Olaparib), 100 mg/kg intraperitoneally, three times per week. 3: AT1 group, which was treated with PARPi (Olaparib) by the AT1 algorithm (dose modulation; see below). 3: AT2 group, which was treated with PARPi (Olaparib) by the AT2 algorithm (dose skipping; see below). Tumor growth was monitored every other day and tumor size was measured by calipers three times a week (Monday, Wednesday, Friday). These measurements were used to inform the dose choices under AT1 and AT2 at these times. Tumor volume was calculated using the following formula: volume = π (short diameter)^2^ × (long diameter)/6. When the tumor volume reached 200 mm^3^, treatment was started. Animal weights were measured and recorded twice weekly, and the overall health of each animal was noted to ensure timely end points within the experiment. Finally, animals were humanely killed and tumors were extracted.

#### Adaptive therapy with dose modulation (AT1)

Given the observed delay in treatment response, all animals were initially treated every other day for at least 5 days before dose modulation was started (100mg/kg). As soon as the tumor stopped growing the subsequent treatment dose was reduced to 50% of the original dose. When the tumor started growing again (any measurable growth from the previous time point), we applied the original dose again and whereas if the tumor stayed under control we reduced the dose by another 50%.

#### Adaptive therapy with treatment skipping (AT2)

Treatment started at MTD (100 mg/kg) for at least 5 days, and subsequently continued until the tumor stopped growing (no measurable growth from the previous time point). As soon as tumor size growth stopped or reduced, we skipped the next treatment. Treatment started again as soon as the tumor started growing as measured by caliper.

### 2.6. Mathematical model development

The aim of our mathematical model was to describe and predict the tumor population size over time in response to different treatment schedules (as measured by the percentage of the well covered by cells, which we will refer to as *confluence*). Given the continuous nature of the confluence measurements, we chose to model the population dynamics using ordinary differential equations (ODEs), such that *N (t)* (in % confluence) represents the confluence at time t (in days). We developed our model in three consecutive steps (Figure 2): Firstly, we identified terms to describe untreated growth (Figure 3). Next, we characterized the dynamics in response to continuous treatment at 100µM Olaparib (Figure 4), and lastly, we extended this model to cover the response at several different drug doses: 10, 50 and 100µM (Figure 5). At each step, we compared different plausible models and picked one to carry forward to the next step. This not only increased confidence in our final model choice, but also was an important step to help elucidate underlying biology by ruling out hypotheses that were inconsistent with the data. Below, we will provide an overview of each model we examined and define the associated model parameters (see Table 1 for an overview). Further justification and interpretation are provided in Section 3.

**Figure 4.**
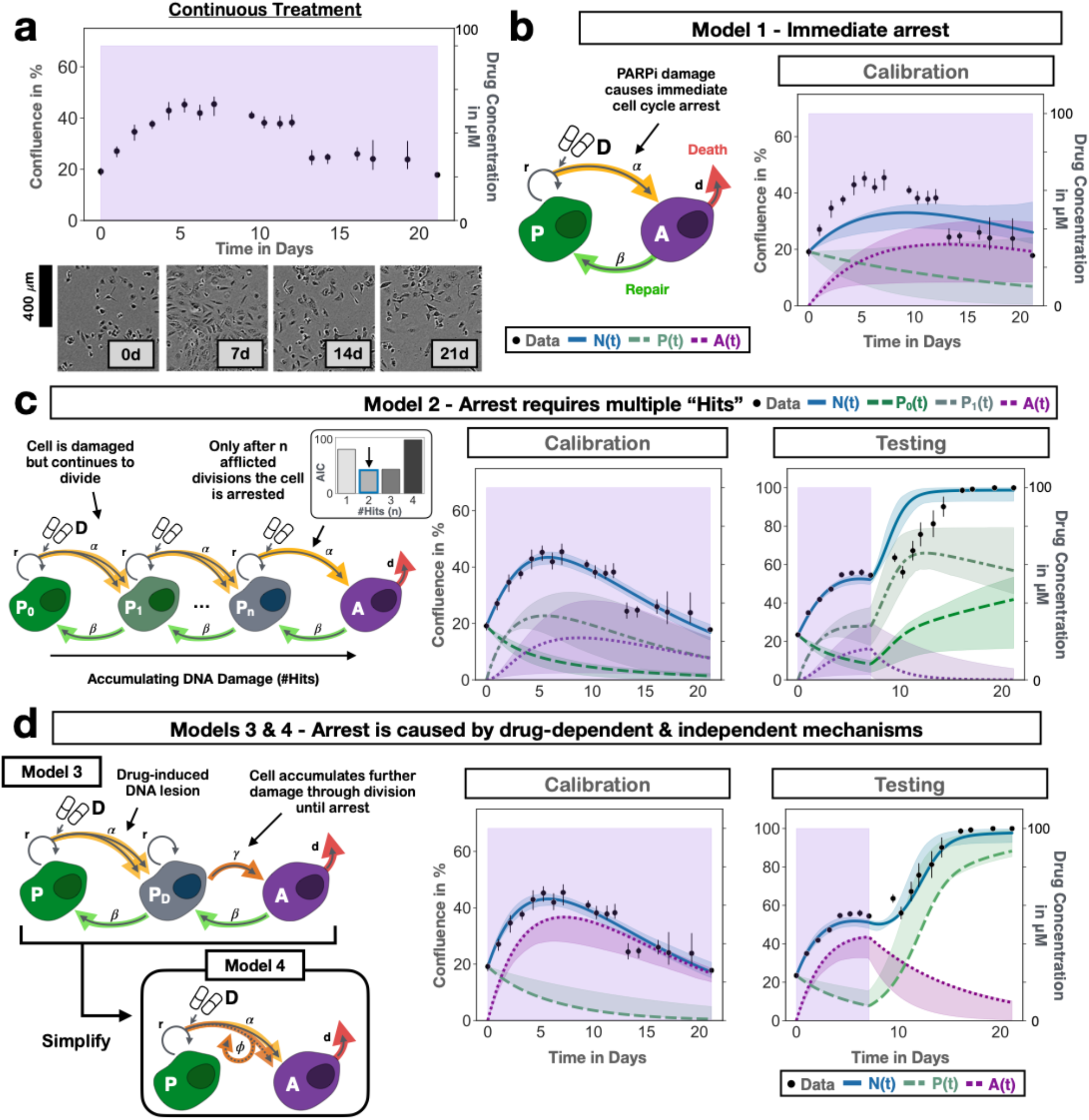
Mathematical modeling of the PARPi treatment response indicates that cells can undergo multiple divisions before entering PARPi-induced cell cycle arrest. Points and bars denote mean and 95% CIs of observed confluence (n=3 independent replicates). Solid lines show the model predictions based on the maximum likelihood estimate, and bands indicate 95% CIs derived via parametric bootstrapping. **a)** Treatment response dynamics measured with our in vitro imaging setup, showing a delayed response where the population initially expands under treatment before it contracts. **b)** A simple model which assumes PARPis induce cell cycle arrest immediately once a cell attempts to divide cannot explain this initial expansion seen in the data (Model 1; Equations (1)-(4)). **c)** A more complex model in which cells need to acquire PARPi-induced damage over multiple rounds of cell division can explain the dynamics under continuous treatment (Model 2; Equations (5)-(9)). Specifically, a value of 2-3 divisions before arrest appears to be most consistent with the data. However, this model predicts faster recovery upon drug withdrawal than what is seen in vitro, suggesting further refinement is required. **d)** To address this, we tested a model which assumes that the DNA damage that results in cycle arrest is initially induced by PARPis but is subsequently exacerbated through cell division independent of further drug exposure (Model 3; Equations (10)-(13)). This can explain the dynamics in response to both continuous and intermittent schedules (for corresponding fits/predictions see Supplementary Figure S4a). Assuming that cells rarely recover from arrest, we were able to simplify this model whilst maintaining high fitting and prediction accuracy, which yielded the final treatment model which we carried forward for our study of treatment scheduling (Model 4; Equations (14)-(16); fits/predictions as shown).

**Figure 5.**
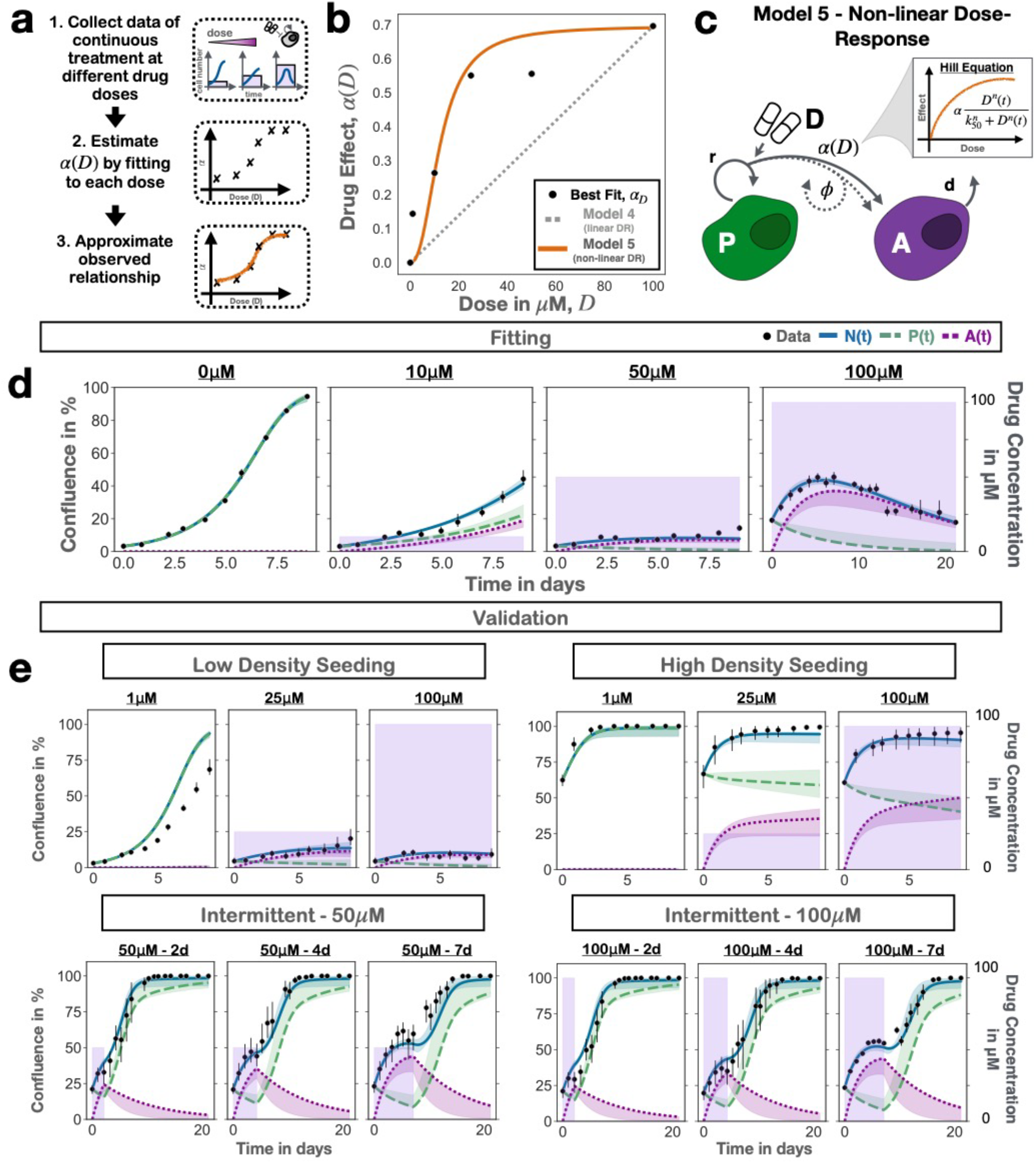
Characterizing the relationship between the treatment dynamics and the drug dose. Points and bars denote mean and 95% CIs of observed confluence (n=3 independent replicates). Solid lines show the model predictions based on the maximum likelihood estimate, and bands indicate 95% CIs derived via parametric bootstrapping. **a)** Approach used to deduce the dose-response relationship. **b)** Empirical dose-response relationship derived from our data, demonstrating a concave curvature, which cannot be described by the linear dose-response model assumed in Model 4 and motivated the illustrated Hill function model (Model 5). **c)** Diagram showing how we extended Model 4 by assuming that the damage probability, α(D), increases non-linearly with dose according to a Hill equation (Model 5; Equations (14)-(17)). **d)** Model fits obtained when calibrating Model 5 with data from 0, 10, 50 and 100µM Olaparib. **e)** Testing of Model 5 on data from 12 different experimental conditions. Together, Panels d & e show that Model 5 can fit and predict the PARPi response of OVCAR3 cells in vitro under various conditions with high accuracy, including the fact that the treatment response varies with cell density.

**Table 1.** Overview of model variables and parameters.

| Variable/<br>Parameter | Description | Range |
| --- | --- | --- |
| $N(t)$ | Tumor population size at $t$ days (in % confluence) | 0-100 |
| $P(t)$ | Size of actively proliferating population at $t$ days (in % confluence) | 0-100 |
| $P_i(t)$ | Size of proliferating population with $i$ rounds of divisions impacted by PARPi-induced damage at $t$ days (in % confluence) | 0-100 |
| $P_D(t)$ | Size of proliferating population with (any) PARPi-induced damage at $t$ days (in % confluence) | 0-100 |
| $A(t)$ | Size of arrested population at $t$ days (in % confluence) | 0-100 |
| $D(t)$ | Drug concentration in the well at $t$ days (in $\mu\text{M}$) | 0-100 |
| $D_{Max}$ | Maximum administered drug concentration (in $\mu\text{M}$) | 100 (fixed) |
| $r$ | Cell growth rate (in $\text{day}^{-1}$) | $10^{-4}$-2 |
| $K$ | Carrying capacity (in % confluence) | 0-100 |
| $v$ | Shape parameter determining curvature of density-dependence relationship (dimensionless) | 0-5 |
| $\alpha$ | Probability of PARPi-induced cell cycle arrest during cell division when treated at $D_{Max}$ (dimensionless) | 0-1 |
| $\beta$ | Rate at which arrested cells repair themselves (in $\text{day}^{-1}$) | 0-0.5 |
| $d$ | Rate at which arrested cells undergo apoptosis and detach from the plate (in $\text{day}^{-1}$) | 0-4 |
| $r_i, \alpha_i, \beta_i$ | Growth rate, damage probability, and repair rate of sub-population with $i$ rounds of PARP-afflicted cell divisions (with units, respectively, $\text{day}^{-1}$, dimensionless, $\text{day}^{-1}$) | |
| $\gamma$ | Rate at which damaged cells become arrested (in $\text{day}^{-1}$) | 0-4 |
| $\phi$ | Average number of divisions a damaged cell undergoes before apoptosis (dimensionless) | 0-2 |

#### Growth Models

We tested 5 different, commonly-used models of untreated tumor growth representative of different assumptions about the strength of density-dependence (see also Figure 3b): i) Exponential growth, which assumes no change in per-capita growth rate with increasing density 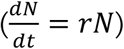, ii) von Bertalanffy growth^44^, which assumes cells grow as a sphere with only the cells on the surface dividing so that the growth rate scales approximately with the sphere’s surface area 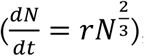, iii) Logistic growth, which assumes a linear decrease in per-capita growth rate with density 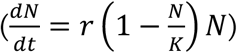, iv) Gompertzian growth^45^, which assumes an exponentially decreasing relationship with density 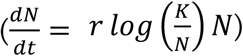, and v) Generalized Logistic growth, which assumes the per-capita growth rate decays according to a power-law 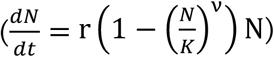. Throughout, *r* (in day^-1^) denotes the instantaneous growth rate, *K*(in % confluence) is the population’s carrying capacity, and *v* is a dimensionless shape parameter.

#### Treatment Models

We assumed that the Olaparib concentration, *D(t)* (in µM), was homogeneous within a well and due to the regular medium replenishment could be assumed to be piece-wise constant over time.

Since it takes time for cells to undergo apoptosis and detach from the plate, we divided the population into an unaffected, proliferating compartment, *P(t)*, and an affected, arrested compartment, *A(t)*. To investigate and characterize the treatment response, we examined 4 treatment models with different assumptions about the conditions required for a cell to be forced into apoptosis (note that for each model, we assume generalized logistic growth, as this was the model selected from the 5 growth models we considered):

- ***Model 1:*** During mitosis, cells in the *P* compartment have a probability *α(D)* (dimensionless) to acquire PARPi-induced DNA damage, which is a function of the PARPi concentration, *D(t)*. If this happens, then division is immediately aborted, the cell becomes arrested and undergoes apoptosis at rate *d* (in day^-1^), unless it is able to repair itself, which occurs at rate *β* (in day^-1^):

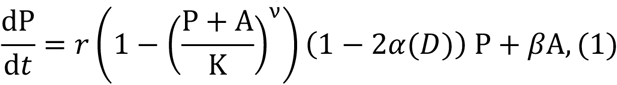

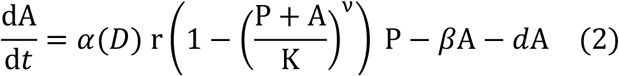

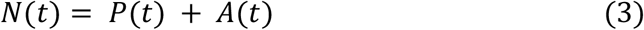

where for simplicity we initially assumed in Models 1-4 that the relationship between drug concentration and damage probability was linear:

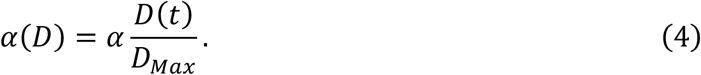 Here, *D*_*max*_= 100µM is the maximum administered drug concentration, introduced for scaling purposes, and *α* is the damage probability (dimensionless) when treated at *D*_*max*_. To represent the fact that cells are most sensitive to PARPi-induced damage when they are undergoing mitosis, we assumed in Equations (1) & (2) that the rate at which cells are arrested by treatment is proportional to the population’s growth rate. The factor of 2 in Equation (1) accounts for the fact that due to the arrested division no daughter cell will be produced.
- ***Model 2:*** This model assumed that a cell needs to acquire multiple PARPi-induced lesions before arrest is induced, so that the population is composed of *n* different proliferating sub-populations with increasing levels of DNA damage, denoted by *P*_*i*_*(t)* (for *i*= 0, …,*n*; in units of % confluence for all compartments):

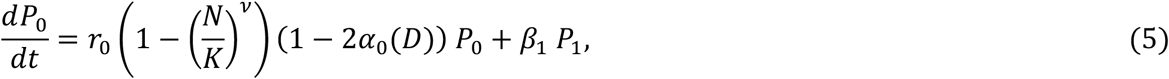

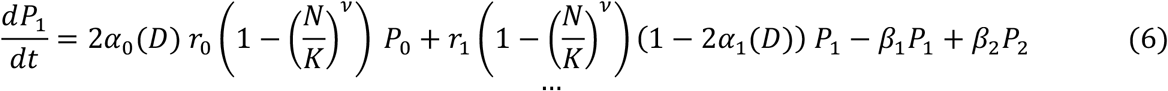

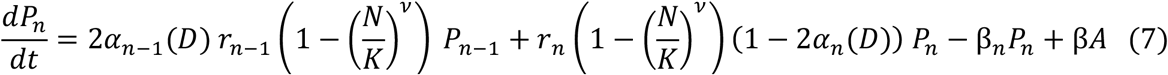

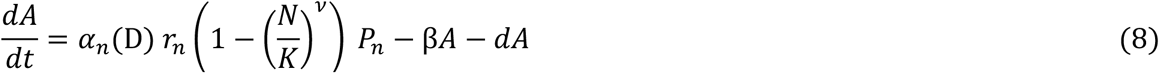

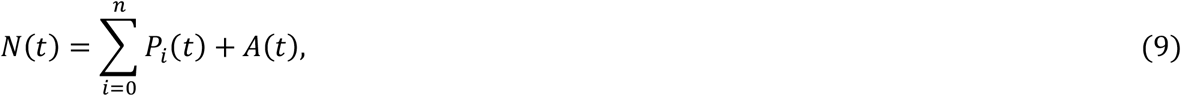

where *r*_*i*_, 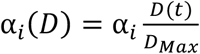, and *β*_*i*_ are the growth rate, concentration-dependent probability of drug-induced damage, and repair rate for sub-population *P*_*n*_, respectively. For the analyses presented in the main text of the paper, we assumed that DNA damage did not change the characteristics of the cell until *n* rounds of damage had been acquired (i.e. *r*_*i*_ = *r, α*_*i*_ = *α*, and *β*_*i*_ = *β* for all *i*= 0, …,*n*). Additional analyses where we allowed *r*_*i*_ and *α*_*i*_ to vary are shown in Supplementary Figures S2-S4.
- ***Model 3:*** While in Model 2 continued drug exposure was required for a cell to keep accumulating DNA damage, this model assumed that the presence of drug was only necessary for induction of an initial lesion. Damaged cells, *P*_*D*_*(t)*(in % confluence), might then continue to divide but would become arrested at a drug-independent rate γ (in day^-1^):

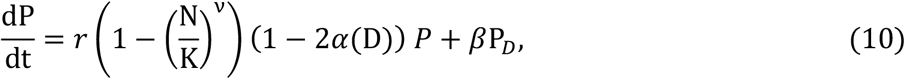

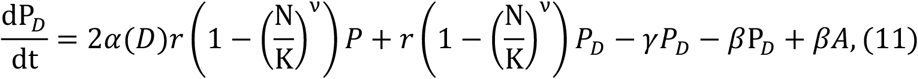

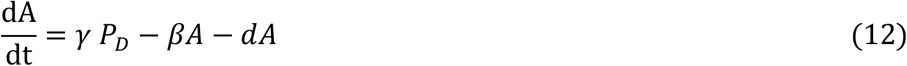

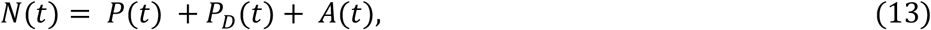

where again we assumed a linear dose-effect relationship (*α*(*D*) given by Equation (4)).
- ***Model 4:*** This model was a simplified version of Model 3, derived by assuming that repair was negligible. In addition, this model assumed that the extra divisions a damaged cell may undergo before arrest could be summarized in a single step, so that a cell that was damaged by drug would on average give rise to ϕ (dimensionless) arrested daughter cells:

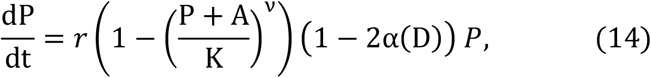

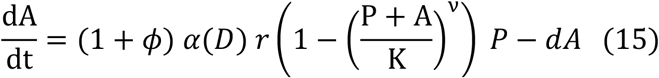

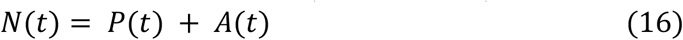 Like Models 1-3 this model assumed that *α*(*D*) was linear (Equation (4)).

#### Dose-Response Model

In the final step, we explored the relationship between dose and treatment effect. We extended Model 4 by assuming that the drug effect was non-linear, so that the damage probability in Equations (14) & (15) was given by:

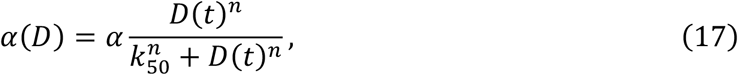

where *k*_50_ (in µM) was the drug concentration at which half the maximum possible effect was achieved and *n* was a non-dimensional shape parameter. This gave us the final model we carried forward for prediction-making, referred to as Model 5 (Equations (14)-(17)).

### 2.7. Model calibration and validation

We calibrated each model by using a Maximum Likelihood approach in which we minimized the root mean squared error (RMSE) between the model-predicted confluence, *N*(*t)*, and the experimentally observed data. Specifically, we fitted to the average of the measured confluence across the three replicates per time point. Following the three steps outlined in Figure 2, we used data from three sets of experimental conditions to sequentially infer the parameters related to growth (*r, K*, and *v*), treatment (*α, β, d*,γ, and ϕ), and dose-response components of the model (*n* and *k*_50_), respectively. When transitioning from one step to the next all parameters related to the prior component(s) were kept fixed. When inferring *n* and *k*_50_, we fitted to data from three conditions simultaneously (continuous treatment at 10, 50, and 100µM) by minimizing the combined RMSE across the three conditions. For initial conditions, we assumed that all cells were initially in the proliferating compartment so that *P*(0) (or *P*_0_(0) for Model 2) was equal to the observed confluence at time 0, and all other compartments were set to 0. Initial conditions were not allowed to vary during fitting.

To test the ability of our models to predict the treatment dynamics under unseen experimental conditions, we performed three sets of validation experiments (Figure 2). In these, we set the initial conditions in the ODE model equal to those observed *in vitro* (again assuming all cells to be in the *P* compartment), and compared the dynamics predicted by simulating the model forward with that observed experimentally. Importantly, all parameters were kept fixed in these experiments.

### 2.8. Uncertainty quantification

Parametric bootstrapping was used to estimate the uncertainty in our parameter estimates and model predictions. To do so, we used the fitted model to simulate 250 synthetic experimental replicates 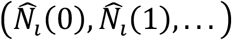 for *i* = 1, …, 250, by sampling residuals from the error model as follows: 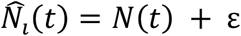, where *ε* ∼ 𝒩(0, σ_*ε*_) is the residual and σ_ε_ is the residual variance of the Maximum Likelihood model fit, 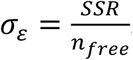. Here, *SSR* is the sum of squared residuals of the Maximum Likelihood fit and *n*_*free*_ is the number of free (fitted) parameters. Next, we fitted the model to each of the synthetic replicates using the same protocol as when fitting the real data. Unless otherwise stated, each of these optimization runs was started from a different random guess within the parameter space. This yielded a distribution of bootstrap estimates for the model parameters and model predictions, from which we derived the presented confidence intervals. To propagate the uncertainty when proceeding from estimating the growth (*r, K*, and *v*), to the treatment (*α, β, d, γ*, and ϕ), and subsequently the dose-response parameters (*n* and *k*_50_), we applied the following protocol: for each bootstrap replicate *i*, we set the fixed parameters to the values obtained in *i*^*th*^ bootstrap during the step in which we estimated these parameters. For example, the value of *r* in the 1^st^ bootstrap for Models 1-5 was taken from the 1^st^ bootstrap when estimating *r* from fitting to the growth data in Step 1 of the model development process. The reason why we chose a parametric method rather than a more assumption-agnostic, non-parametric method was that we only had three replicates available per experimental condition. We also tested uncertainty estimation using the delta-method^46^, which yielded comparable results (not shown).

### 2.9. Numerical methods

All data analyses, model fitting and simulations were carried out in Python 3.8. Specifically, we used the `DOP853` explicit Runge-Kutta method in scipy 1.6.2 to solve the ODEs, and the lmfit package^47^ (version 1.0.2) and the Levenberg-Marquardt algorithm implemented in the least_squares method in scipy to carry out model fitting. Visualizations were produced with Pandas 1.2.4, Matplotlib 3.5.2, and Seaborn 0.11.1. All code is available at: https://github.com/MathOnco/PARPi_Model.

## 3. Results

The aim of this paper was to investigate the feasibility of alternative treatment schedules for Olaparib which maintain or even enhance tumor control but reduce drug use. To do so, we performed a series of *in vitro* experiments to develop, calibrate, and validate a mathematical model of the treatment response dynamics. We proceeded in three steps, gradually complexifying the dosing regimens that our model could describe (Figure 2). In the following, we will report on the insights gained into the treatment dynamics by our modeling and subsequently discuss their implications for scheduling. We repeated our analysis on two commonly used human epithelial ovarian cancer cell lines (OVCAR3 and OVCAR4). These were originally derived from ascites and were chosen to model experimentally the peritoneal disease that PARPi maintenance seeks to manage. As both cell lines yielded similar results, we focus on OVCAR3 and refer to the supplementary material for the OVCAR4 results.

### 3.1. Untreated cells growth dynamics indicates non-linear density dependence

The cytotoxic effect of PARPis is based on their interference with the DNA replication machinery and the induction of double strand DNA breaks. To begin with we therefore analyzed the growth dynamics of OVCAR3 cells in the absence of treatment in order to accurately capture the fraction of dividing cells over time. Using time-lapse microscopy and image analysis, we collected images daily, such as those shown in Figure 3a, and from these quantified the population size over time as a percentage of how much of the visible area was overgrown (% confluence). Subsequently, we compared the observed trajectory with five plausible and commonly used mathematical models, representing different assumptions about how rapidly the fraction of dividing cells decreased as the population approached confluence (their “density dependence”; Figure 3b; see Section 2.6. for the equations). We found that a generalized logistic model was the most consistent with the observed growth dynamics, even when we penalized for its additional parameters (Figure 3b). The corresponding concave shape of the density dependence relationship suggests reduced contact inhibition, consistent with the cancerous nature of these cells. Repeating this analysis with OVCAR4 cells showed stronger density dependence but yielded otherwise similar conclusions (Supplementary Figures S1a & b).

Next, we sought to test how well this model could predict growth under experimental conditions different to those for which we had calibrated it. Given that AT relies on cells competing in close proximity^33,37,40^, we chose to test the model in its ability to predict growth when we seeded cells at a higher initial density (60% confluence). This analysis corroborated our choice of the generalized logistic model, although for both cell lines it slightly under-predicted the initial growth rate of the population (Figure 3c & Supplementary Figure S1c). The parameter estimates for each cell line are summarized in Supplementary Figures S1d & e, respectively.

### 3.2. Even at high doses, there is a time delay before the population begins to shrink

Having characterized the cells’ growth dynamics, we turned to study their response to therapy. To do so, we exposed the cells to continuous treatment at 100µM Olaparib for 21d. This revealed that, despite high-dose treatment, the population tended to initially expand before a treatment-induced regression could be observed (Figure 4a). To understand the reasons behind this delay in treatment response, and to develop a means for subsequent *in silico* schedule optimization, we next extended our mathematical model to capture these dynamics.

PARPis interfere with the DNA repair and replication machinery which induces cell cycle arrest, and eventually results in apoptosis if cells are unable to repair themselves. Based on this understanding, we tested a model in which we assumed that the tumor population could be divided into two subpopulations (Figure 4b): i) Cells which are actively cycling and unaffected by PARPi (*P*), and ii) cells in PARPi-induced cell cycle arrest (*A*). Further, we assumed that during treatment drug caused cell cycle arrest in a fraction of cells in the proliferating subpopulation (those undergoing mitosis during this period) and thereby moved them from the *P* to the *A* compartment. Once arrested, these cells would either repair themselves and return to the proliferating compartment or would undergo apoptosis and detach from the plate. Seeking to keep our model as simple as possible, we initially assumed that if there was Olaparb induced damage, then the cell would immediately abort division and go into arrest (Model 1; Figure 4b; Equations (1)-(4)). Fitting this model to our data, we found that it was able to reproduce the biphasic behavior of expansion and contraction we had observed experimentally (Figure 4b). However, whilst qualitatively in agreement, the model predicted a much less pronounced initial expansion than what we had seen *in vitro*, suggesting that the assumption of immediate cell cycle arrest was inconsistent with our data (Figure 4b).

### 3.3. Modeling indicates that cells undergo 1-2 extra divisions before PARPi-induced arrest

Based on this observation, we tested a model in which we assumed that multiple divisions under PARPi exposure were required to amount sufficient damage to induce cell cycle arrest (Model 2; Equations (5)-(9)). In this model, the cell accumulated DNA damage if affected by the PARPi during mitosis but it still successfully completed cell division. Only after too many “hits” were received, was the cell forced to abort division and was pushed into cell cycle arrest (Figure 4c). In addition, cells could repair damage and for simplicity we assumed that partial damage did not alter the behavior of the cell, so that parameters were the same across all proliferating compartments, *P*_*i*_. Fitting this model, we found that assuming a cell underwent 2-3 cell divisions before it was forced into arrest, reliably reproduced the *in vitro* data (Figure 4c & Supplementary Figure S2a; r^2^ values of 0.91 and 0.9, respectively). We concluded that Olaparib-induced damage did not appear to induce arrest immediately, but that instead cell death resulted from the build-up of further damage over multiple rounds of cell division.

### 3.4. PARPi cytotoxicity involves both drug-dependent and independent steps

Given this observation, an important follow-up question was whether treatment interruptions would interfere with the action of the drug. If cells needed to be damaged not just once, but multiple times, and the continued presence of drug was necessary to induce this damage, then this would mean that withdrawal of treatment too early may not allow for enough time to induce cell death. To investigate this, we simulated an experiment in which we treated cells for varying lengths of time (1d, 2d, 4d, 7d), after which we removed treatment. As expected, the model predicted that the population would start growing again within 24h after drug removal (Figure 4c and Supplementary Figure S2b). However, comparing these predictions with the dynamics when we repeated this experiment *in vitro*, we found that the recovery predicted by the model was too fast (Figure 4c and Supplementary Figure S2b). This suggested that the cells were continuing to experience the impact of the PARPi-induced damage even once treatment had been withdrawn.

To characterize the way in which the cells were impacted, we tested whether this lingering effect took the form of either a decreasing proliferation rate or an increasing drug sensitivity as damage accumulated in a cell, but neither model was able to explain the data (Supplementary Figures S2b & S3a-b). Examining why this was the case, we found that while decreasing the proliferation rate reduced the growth rate, it also reduced the cells’ drug sensitivity, resulting in a build-up of damaged, but still proliferating, cells which explained why the model predicted too fast a regrowth upon drug withdrawal (compare the levels of P_1_(t) in Supplementary Figure S2b). This prompted us to revisit our assumption that drug exposure was required for the further build-up of DNA damage after an initial PARPi-inflicted lesion. So, we iterated testing with a refined model in which cells continued to divide after PARPi damage but eventually underwent apoptosis independent of further treatment, unless they had been able to repair themselves (Model 3; Figure 4d; Equations (10)-(13)). This model was able to recapitulate the treatment response under continuous, as well as intermittent, treatment with high accuracy, suggesting that PARPi-induced cytotoxicity involved both drug-dependent and independent steps, as might be expected from PARP trapping. (Supplementary Figures S2b & S3a-b). Repeating these analyses with the OVCAR4 cells corroborated this result (Supplementary Figure S4a-c) and adds support to the growing evidence for the importance of PARP trapping in PARPi action.

### 3.5. A slow repair rate simplifies the model needed for prediction-making

Having investigated how PARPis damage cells, we turned to consider the question of the rate of repair. Examining the estimates for *β* provided by Models 1-3 for both cell lines indicated that little repair appeared to be taking place (Supplementary Figures S3c & S4d). This observation not only provided further biological insight, but also suggested a way of simplifying our model. While Model 3 was useful for gaining a mechanistic understanding of the actions of Olaparib, its complexity meant that it was difficult to parameterize it with the data at hand, seen, for example, in the significant uncertainty associated with the sizes of the individual subpopulations (Supplementary Figure S2b). By neglecting repair, we were able to reduce our model back to two populations, consisting of healthy proliferating cells, *P(t)*, and arrested cells on the way to apoptosis, *A(t)*. In this way, the transient rounds of cell division following PARPi-induced damage could be combined into a single step, where a new parameter ϕ captured the number of divisions a damaged cell would undergo before cell cycle arrest (Model 4; Figure 4d; Equations (14)-(16)). For both cell lines, this model provided fits and predictions as good as, if not better than, the more complex Model 3, with less uncertainty in its predictions and parameter estimates (Figure 4d & Supplementary Figures S3 & S4).

### 3.6. Analysis of the response at different doses reveals positive cooperativity in drug action

In the last step of model development, we sought to characterize how the treatment dynamics varied with drug dose (Figure 2). This was so that we could subsequently use the model to investigate treatment algorithms which adapted not just whether or not treatment was given, but also adjusted the dose. To do so, we first used our *in vitro* time-lapse imaging pipeline to measure the response dynamics of cells continuously exposed to 1, 10, 25, and 50µM of Olaparib for 9 days. We then fitted Model 4 to each drug level, allowing the treatment-induced damage probability, *α*(*D*), to vary freely with dose (Figure 5a). This revealed a concave dose-response relationship for *α*(*D*), indicating that the dose relationship was not linear, as we had assumed in Model 4, but that there was evidence for positive cooperativity in PARPi action (Figure 5b). This meant that acquiring a PARPi-induced lesion appeared to increase the probability that a cell would suffer further PARPi-induced damage. We also explored whether ϕ or *d* varied with dose but did not find evidence to support this (Supplementary Figures S5a & b).

To integrate this positive cooperativity into our mathematical model, we extended Model 4 by introducing a Hill function to describe the relationship between the dose and the treatment-induced damage probability, *α*(*D*) (Model 5; Figure 5c; Equations (14)-(17)). After calibrating the shape parameter, *n*, and half-effect parameter, *k*_50_, using the data at dose levels 10, 50 and 100µM we found that this model was able to closely recapitulate the experimentally observed drug-response relationship (Figure 5b), as well as the associated treatment dynamics (Figure 5d; see Figure S5c & d for a summary of the parameter estimates). Repeating this analysis with a different set of “training” doses (e.g. 1, 25, 100µM; not shown) and with OVCAR4 (Supplementary Figures S6a & b) corroborated our conclusions.

### 3.6. The model is highly predictive and reveals that drug response changes with cell density

To validate the final form (Model 5) of our mathematical model, we tested its ability to predict the treatment dynamics under combinations of different conditions (varying doses, seeding densities, and continuous vs intermittent schedules). We found that for both cell lines our model was able to predict the observed dynamics with high accuracy (Figure 5e and Figures S5d & S6c-f; see Figure S6g for the OVCAR4 parameters). In particular, our mathematical model predicted that the cells would recover quickly after drug withdrawal and would experience a certain protection from treatment when grown at higher density. These predictions were validated *in vitro* (Figure 5e) and suggested that how, and when, treatment was adapted would have to be carefully planned. Thus, in the final part of this study, we leveraged our calibrated and validated mathematical model to study different possible adaptive treatment strategies.

### 3.7. Model simulations of different adaptive therapy algorithms indicate dose modulation may perform better than dose skipping

In the prostate cancer adaptive therapy trial by Zhang, et al. ^42^, the authors alternated between drug administration and drug holidays in order to keep tumor size between the baseline value at the start of treatment and 50% of this value (as measured by PSA). However, PARPi maintenance therapy immediately follows systemic therapy and possibly surgery or radiation therapy, so that there is typically little or no evidence of remaining disease at the start of treatment^14,17^, making a strategy similar to that of Zhang, et al. ^42^ difficult to implement. As an alternative, we investigated two previously published adaptive algorithms which adjust treatment based, not on tumor size, but on *changes* in size (Figure 6a): i) AT1^1,32^, which modulates the dose administered at the current time point, increasing it if the tumor grows too quickly and decreasing it if tumor growth slows sufficiently, and ii) AT2^1^, which performs dose-skipping akin to Zhang, et al. ^42^, except that doses are skipped when the *growth rate* drops below some threshold. To make these algorithms easier to implement we made two simplifications compared to Enriquez-Navas et al^1^: for AT1 we selected from one of only five dose levels (0, 12.5, 25, 50, 100µM), separated by factors of *α* = 2, and for AT2 we assessed growth rate over one-step rather than two-step intervals (Figure 6a).

**Figure 6.**
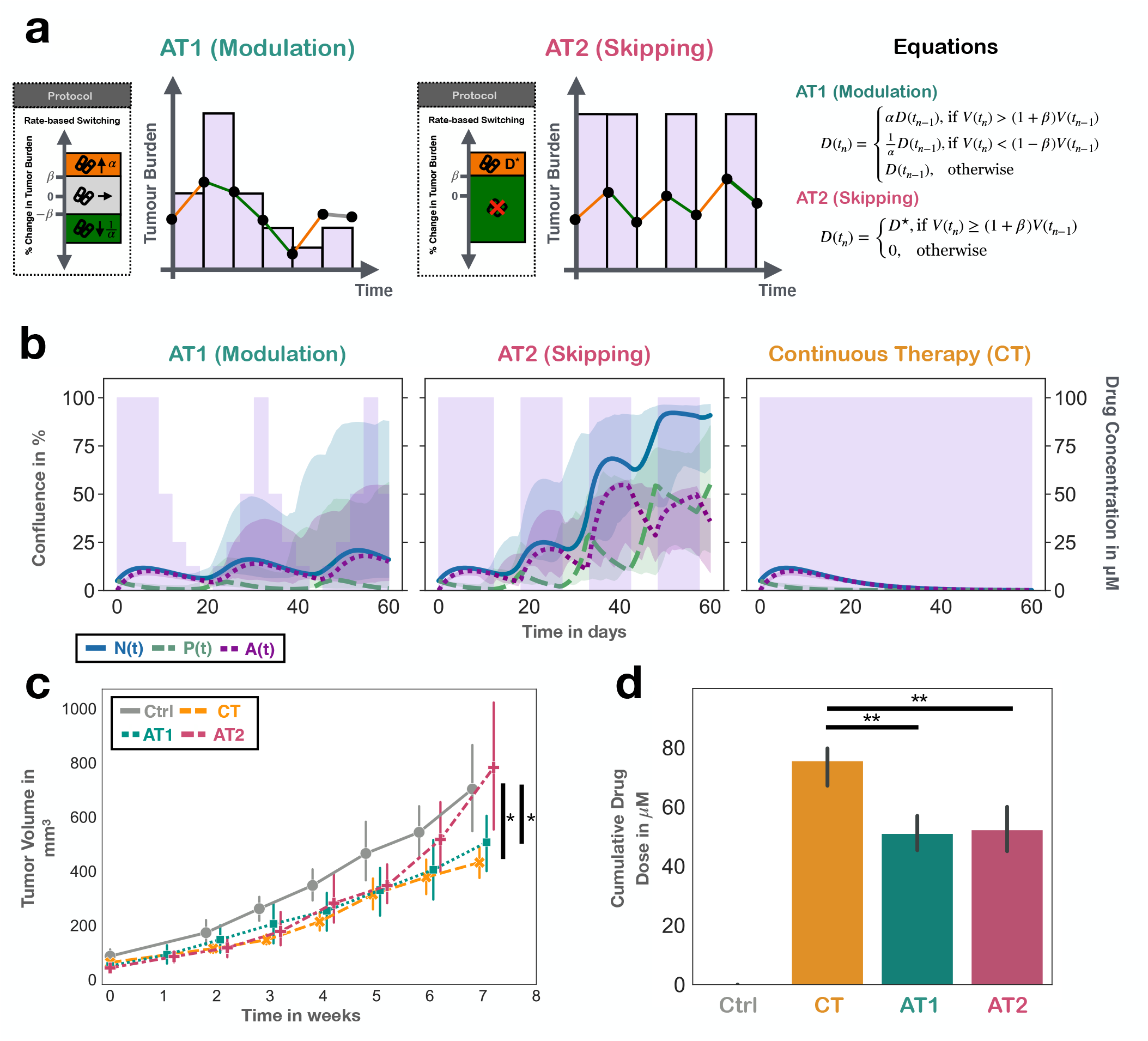
Mathematical modeling and in vivo experiments suggest that adaptive treatment strategies which modulate dose but not completely withdraw it are superior to algorithms based on treatment skipping. Points and bars denote mean and 95% CIs of observed confluence. Solid lines show the model predictions based on the maximum likelihood estimate, and bands indicate 95% CIs derived via parametric bootstrapping. **a)** Two candidate adaptive therapy algorithms, modified from Enriquez-Navas, et al. ^1^ used as starting points for the development of a PARPi-specific strategy. Both adjust treatment based on the tumor’s growth rate, but differ in how these adjustments are made: AT1 modulates the dose, whereas AT2 will completely skip treatment. **b)** Model simulations of both adaptive algorithms as well as continuous therapy using Model 5, showing that care needs to be taken in how treatment is adjusted, with modulation appearing to be superior to dose skipping. The plotted treatment schedule (pink bars) corresponds to the prediction made using the maximum likelihood fit (lines). Tumor parameters: n_0_ = 5%, f_A_ = 0%; AT1: *α* = 2, *β* = 0, D(0) = 100µM; AT2: D^⋆^ = 100µM, *β* = 0, D(0) = 100µM. **c)** & **d)** Tumor growth data (c) and cumulative drug usage (d) in in vivo experiments comparing the three strategies from Panel b, confirming the model predictions (n=6 animals per group). Note: in c), for readability we separated different treatment arms with a small offset along the x-axis, but all measurements were taken at the same time. t-test statistics: * p<0.05, **:p<0.01.

Using our mathematical model, we simulated both algorithms and found that, as anticipated from Section 3.6, there were stark differences in outcome between the two strategies (Figure 6b). AT1 was predicted to be able to maintain the population size relatively small throughout treatment whereas under AT2 the tumor was predicted to rapidly expand. The reason for this were the treatment breaks during AT2, which occurred earlier and more often than under AT1. Conversely, continuous treatment at 100uM was predicted to eradicate the population (Figure 6b). This is because during treatment breaks the surviving proliferating cells in the simulations would rapidly recover, and the delayed treatment response meant that a prolonged drug exposure was required to shrink the population size again. In addition, for both AT1 and AT2 we found that the model predicted considerable variability in the possible trajectories under adaptive therapy, indicating that even small differences in tumor parameters, or treatment timing, can potentially result in distinct outcomes (Figure 6b). Together, these are important observations as these treatment failures occurred despite the absence of drug resistance in our model, implying that how we adapt treatment in this setting has to be carefully planned.

To test these predictions empirically, we attempted to repeat these experiments *in vitro*, but found that it was not possible to culture cells for multiple treatment cycles in the same dish without replating (not shown). Thus, we moved to an *in vivo* setting which provided confirmatory evidence that reducing the cumulative drug use is possible but should be done via dose modulation rather than treatment interruptions (Figures 6c & d; for individual trajectories, see Figure S7; note: to simplify experiments, we only used four dose levels for AT1: 0, 25, 50, 100µM).

## 4. Discussion

PARPis are promising to revolutionize ovarian cancer care, but toxicity, financial costs, and drug resistance mean that not all patients benefit equally, and often improvements are only temporary. Recent results in androgen-deprivation treatment of prostate cancer have shown that by adapting treatment to the treatment response of the individual patient, it may be possible to delay progression and reduce drug use^42,43^. In this paper we took a first step towards exploring adaptive therapy for PARPis by developing a mathematical model with which we could, relatively cheaply, test different plausible adaptive algorithms before proceeding to more expensive *in vivo* experiments.

To the best of our knowledge, our model is the first mechanistic mathematical model of PARPi treatment in the literature, and it was systematically derived from, and validated with, *in vitro* experimental data. Through the process of developing and analyzing our model we made three key observations about PARPi treatment dynamics. Together these indicate that it may be possible to reduce treatment whilst maintaining control over the tumor, but how treatment is adapted will need to be carefully planned.

Firstly, there was a delay in the drug response, which meant that even when treated continuously at a relatively high dose of 100µM (the physiological dose is around 20µM) it would take around 7 days before the population would begin to recede. Importantly, during this time the cells continued to expand, so that even once the population began to shrink it would still take further time on treatment until the total population size dropped below its starting size. Through our integrated modelling approach, we systematically evaluated different plausible mechanistic explanations for this delay, which suggested that despite acquiring PARPi-induced DNA damage, cells underwent 1-2 further rounds of cell division before cell cycle arrest was induced. This is consistent with a recent study by Liu, et al. ^48^ who made similar observations for the dynamics following radiation-induced DNA damage. At the same time, we found that when treatment was withdrawn, the cells would readily recover, even if the population had previously begun to recede under treatment. Together, we draw two conclusions for possible adaptive PARPi therapies: i) drug vacations (if any) will likely have to be shorter than drug on-times, and ii) rather than re-evaluating the patient at regular intervals, it may be beneficial to perform more frequent follow-up during off-treatment periods to account for the asymmetry in the response dynamics.

Secondly, when comparing the response dynamics at different drug doses we found that there was a strong non-linear relationship between dose and drug effect for both cell lines. The concave nature of this relationship implies a diminishing return of increasing the dose. Crucially, this is not only consistent with the wide therapeutic window of Olaparib observed clinically^11,49,50^, but it also implies that schedules with large changes in dose, such as the dose-skipping AT2 algorithm, will have a smaller average rate of tumor suppression than those regimens which maintain a more constant dose, such as the dose-modulating AT1 strategy (see also the concept of fragile vs anti-fragile tumors^51^). This was demonstrated by our simulations showing that there was a risk that AT2 could lose control over the tumor, even in the absence of drug resistance. This conclusion was further confirmed by our *in vivo* experiments and suggests that we would expect greater success from dose-modulating rather than skipping-based adaptive PARPi strategies.

Finally, we observed that more densely seeded cells appeared to experience protection from treatment. In support of this observation, others^52^ have found that spheroid cultures are more resilient to Olaparib treatment than 2-D cell culture, and we hypothesize that this is due to fewer opportunities for cell division in denser cultures. According to our simulations, this reduced sensitivity as the tumor expands during treatment breaks was another reason for failure under the dose-skipping, AT2 algorithm.

Based on these results we propose that future work should explore adaptive PARPi treatment protocols which modulate dose, but unlike the protocol used in prostate cancer^42,43^, avoid prolonged treatment breaks. In Figure 7 we illustrate a potential candidate algorithm based on the AT1 strategy in which dose is adjusted according to the rate of change in CA125 levels. To make the algorithm more clinically feasible, we have reduced the number of possible doses to just two: a higher “treatment” dose, *D*_Max_, and a lower, but non-zero, “maintenance” dose, *D*_M*i*n_. Such a protocol is appealing for several reasons. Its lower cumulative dose may reduce toxicity, whilst it may also slow resistance through competitive suppression. Furthermore, it is plausible that because the periods of high dose treatment are short, one might be able to administer higher, “bolus” doses than current MTD for *D*_Max_, and thereby target cells which are only partially resistant to treatment^53,54^.

**Figure 7.**
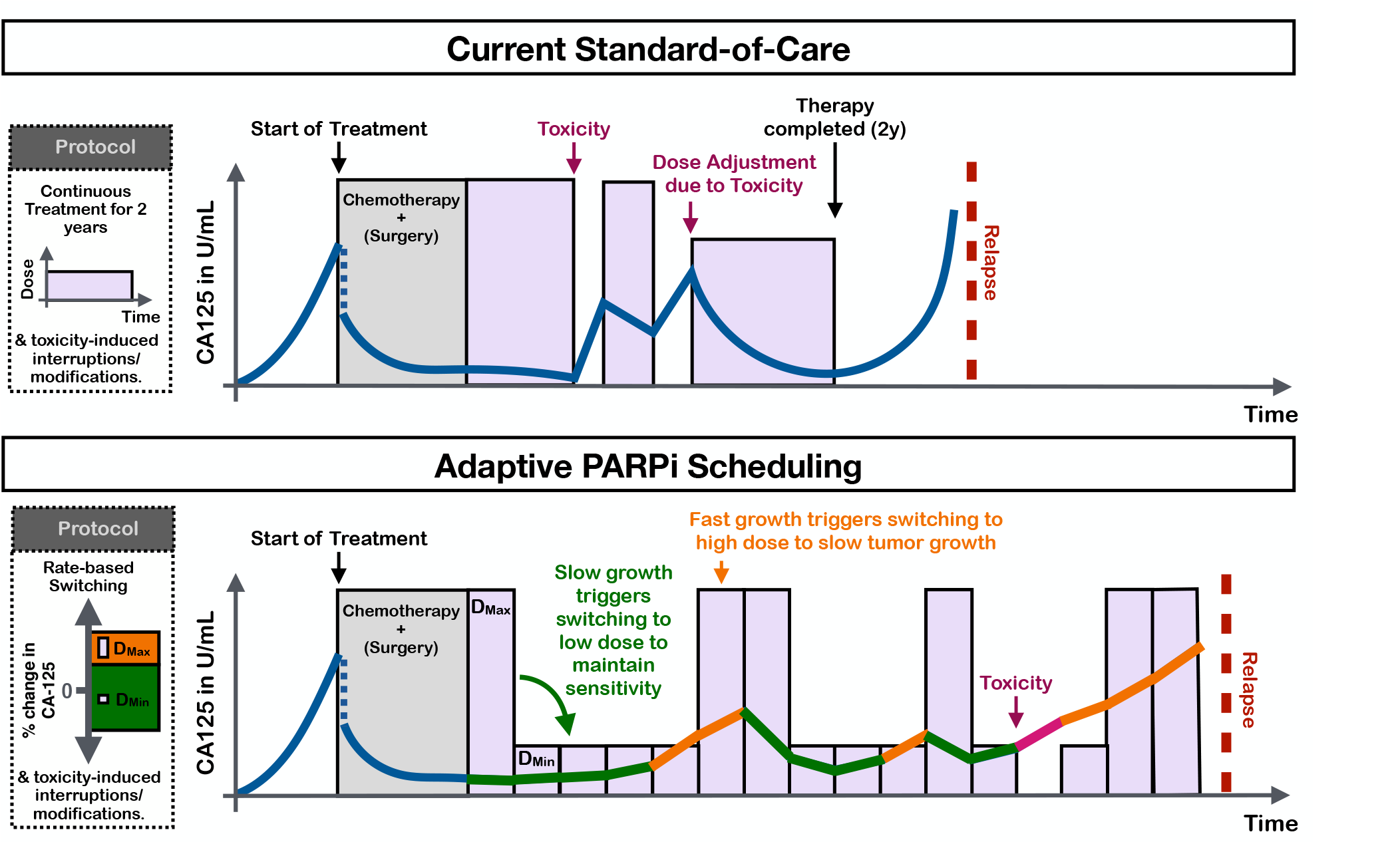
Translational implications of this study. Current standard-of-care administers PARPis continuously for a certain amount of time (2 years in the case of Olaparib) and only adjusts in case of toxicity. In the future, we propose to explore dose-modulating adaptive therapies which, for example, switch between two dose levels depending on the rate of CA-125 level changes. We hypothesize this may reduce toxicity and help to delay relapse by leveraging intra-tumoral competition.

At the same time, our study also highlights potential challenges of adaptive PARPi therapy. Firstly, there is the reduced drug efficacy at higher cell densities. If confirmed in patients, this might limit the maximum tumor burden we could maintain and thereby the competitive suppression we can exert on resistant cells. More generally, our results about the delay in response and the limited scope for holidays indicate that we are likely more constrained in our ability to reduce treatment than has been the case in other adaptive therapy studies so far, where we have typically had, or have assumed, full control over the sensitive cells^36,55,56^. As such, it will be key to extend our work from drug-sensitive to resistant cells and determine whether, or not, our proposed adaptive regimen can delay progression.

To design such an assay, several questions will need to be addressed. Firstly, both cell lines studied in this paper were relatively sensitive to treatment (even if nominally BRCA wild type^57,58^). As a first step, we should therefore test the model on cells with a different HR and/or BRCA status and corresponding drug sensitivity. Secondly, whilst we did partially validate our results *in vivo*, our mathematical model was derived and calibrated by 2-D *in vitro* experiments. There are limitations to such an experimental model: i) growth medium changes to vary drug dose can confound measurements by mechanically disturbing the cells or due to incomplete drug wash-out, ii) cell-cell interactions and nutrient dynamics are limited compared to 3-D, and iii) key elements of the tumor microenvironment are missing, such as endothelial or immune cells. Future research should extend our work to more realistic settings, such as spheroid or mouse models. In addition, the considerable variation in the predicted treatment dynamics depending on the tumor and algorithm parameters in Figure 6, suggests that further work on improving robustness of adaptive therapy is important^41,59^. Finally, it would be useful to experimentally validate the 2-population structure inferred by our mathematical model, for example, by quantifying PARPi-induced DNA damage (e.g. via staining for H2AX or RAD51 foci^60,61^). Backing up the model’s mechanistic underpinnings will help to understand its limitations as we move towards translation.

To summarize, we have presented a systematic analysis of the treatment dynamics during PARPi therapy in ovarian cancer. By closely integrating experiments and mathematical modeling we were able derive insights into the underlying biology and build confidence in our final model. We have intentionally reported the iterative nature of this process, to emphasize that the strength of modeling is not only to rule-in hypotheses (“good fits”) but also to rule them out (“bad fits”). Our work indicates that while there may be scope for adaptive therapy in PARPi treatment of ovarian cancer, careful thought will be required as to how dosing is adjusted (specifically avoiding treatment holidays), and we have proposed a potential adaptive PARPi treatment algorithm to address this. With the growing use of PARPis in other cancers, such as prostate and breast cancer, we believe that our results may be of interest more broadly, and we encourage further exploration of adaptive scheduling as a means for patient-specific toxicity and resistance management in PARPi treatment.

## Acknowledgements

We would like to thank Samantha Byrne and Punit Borad for their assistance with the experiments. The authors gratefully acknowledge funding by the National Cancer Institute via the Cancer Systems Biology Consortium (CSBC) U01CA232382 (supporting M.S., M.R.- T., B. G., A.R.A.A), and support from the Moffitt Center of Excellence for Evolutionary Therapy. This work is partly supported by NCI grants: R01CA249016-01, R01CA272601-01, and U01CA261841-01.

## Author Contributions

M.S., R.W., P.M., M.D. and A.A. conceived and designed the study. M.S. and M.D. collected the experimental data. M.S., J.W., J.G., M.R.T., P.M., and A.A. developed the mathematical model and investigated the implications for adaptive therapy. A.M., R.G. and R.W. provided clinical feedback on the design of the study and the results. M.S. wrote the draft of the manuscript. M.S., J.G., J.W., and A.A. created the figures. All authors subsequently reviewed the manuscript and read and approved the final version. All authors had access to all the data in the study and M.S., P.M., M.D., and A.A. verified the data and had final responsibility for the decision to submit for publication.

## Supplementary Material

**Figure S1.**
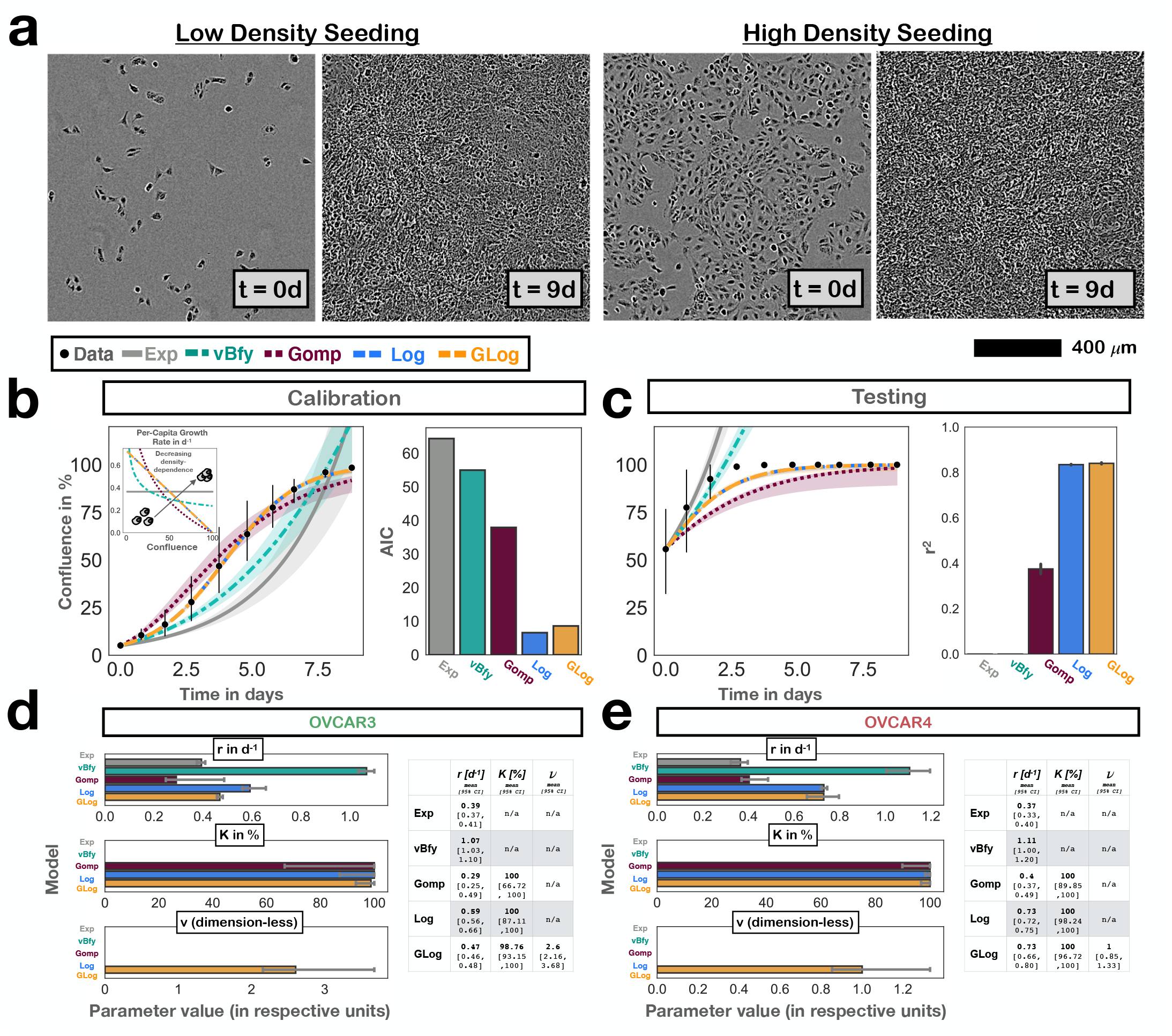
Growth model analysis for the OVCAR4 cells and details of the parameter estimates. In b) & c) points and error bars denote mean and 95% CIs of observed confluence (n=3 independent replicates). Solid lines show the model predictions based on the maximum likelihood estimate, and bands indicate 95% CIs. In d) & e) bars denote the maximum likelihood estimate and error bars show 95% CIs. **a)** Example microscopy images based on which we assessed the growth and treatment dynamics over time. **b)** Results of fitting the 5 growth models to untreated OVCAR4 growth data from cells seeded at low density (Exp: exponential; vBfy: von Bertalanffy; Gomp: Gompertz; Log: Logistic; GLog: Generalized Logistic). In this case the Log model achieved the lowest AIC score. That being said, also the GLog model performed well (ΔAIC = 2), recovering the same linear density-dependence relationship and effectively being reduced to the Log model (v = 1; see also Panel d)). **c)** Testing of the growth models by comparing their predictions for when cells are initially seeded at 60% confluence with the experimentally observed dynamics. As would be expected, the Log and GLog model are equally predictive. Because the GLog model performed very well across both cell lines, we carried it forward as the growth model in our analyses. **d)** Parameter estimates for each model for OVCAR3 cells. **e)** Parameter estimates for each model for OVCAR4 cells.

**Figure S2.**
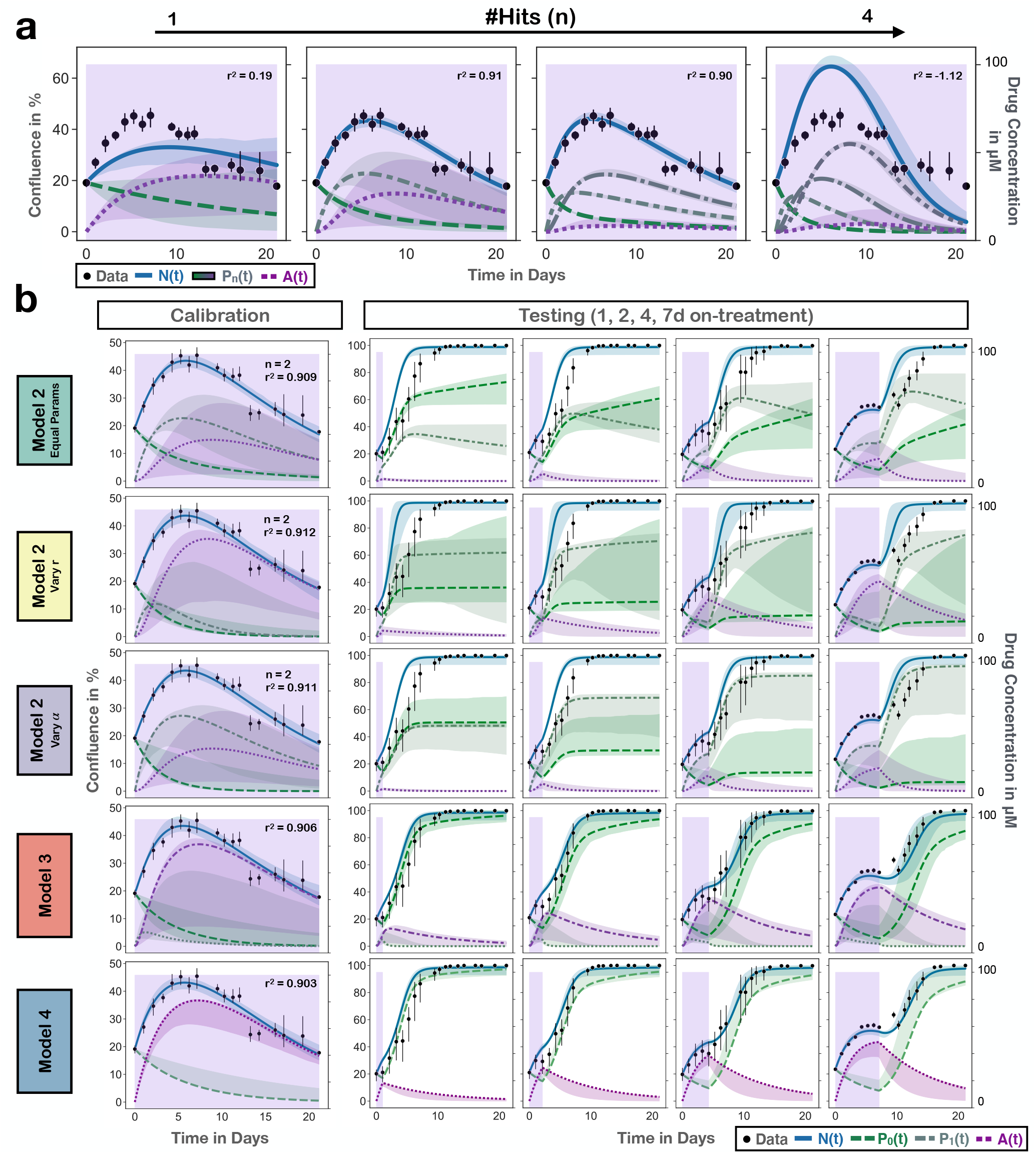
Additional plots documenting the systematic development of the treatment model for OVCAR3 cells. **a)** Model fits of Model 2 for different values of the parameter n, which represents the number of divisions a cell needs to undergo before it accumulates sufficient damage to be pushed into apoptosis. We found a value of n =2 gave the most parsimonious fit (simulations correspond to bar plot in Figure 2c in the main text). **b)** Model fits (left column) and model predictions (right columns) for the five different models examined in order to explain why there is a delay in the treatment response that cannot be accounted for by Model 1. This illustrates how Model 4 provides both the best fit, as well as the most accurate predictions of the testing data, implying that PARPi response involves both drug-dependent and independent steps. Points and error bars denote mean and 95% CIs of observed confluence (3 independent replicates per condition). Lines depict the model predictions based on the maximum likelihood estimate, and bands indicate 95% CIs calculated via parametric bootstrapping. Note that due to issues caused by the existence of local optima in the likelihood surface, we turned off randomization of the initial parameter guesses when fitting Model 2 with varying r.

**Figure S3.**
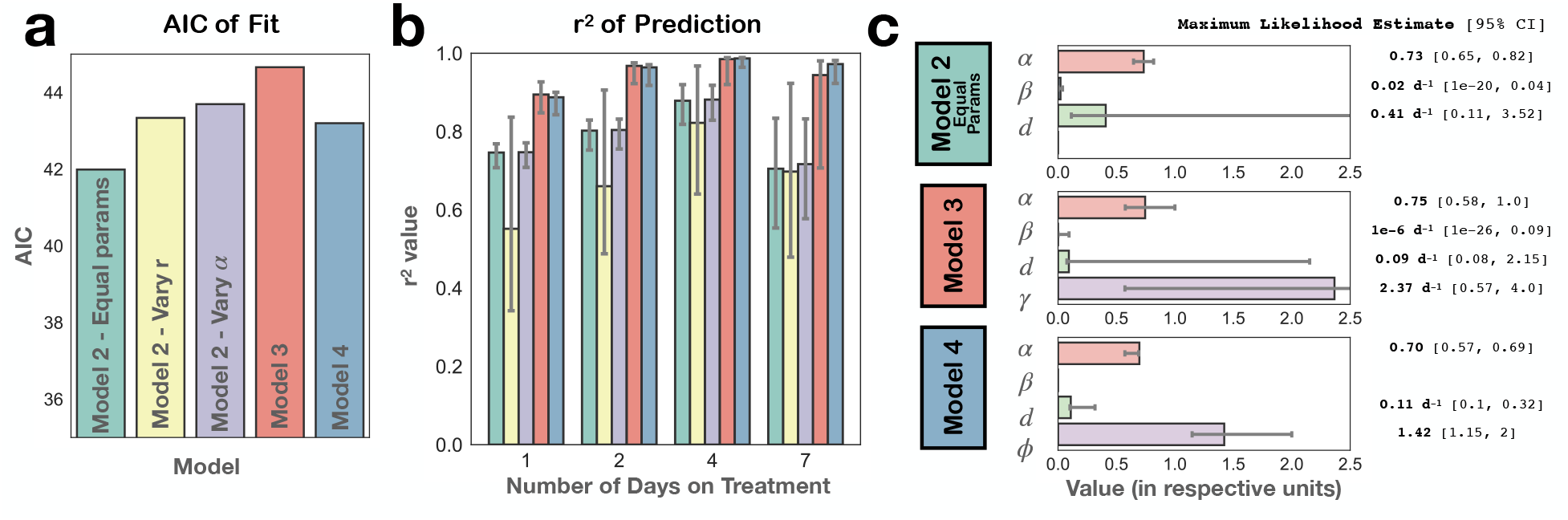
Quantification of the descriptive and predictive power of the different plausible treatment models we explored for the OVCAR3 data, justifying our choice of Model 4 (see Supplementary Figure S2 for the corresponding time dynamics). **a)** Quantification of the goodness-of-fit of each model, showing that all can explain the response under continuous treatment similarly well (most differences in AIC smaller than 2). Thus, we based our choice of which treatment model to carry forward based on their ability to predict the response dynamics for intermittent treatment. **b)** Quantification of the prediction accuracy of each model on the intermittent treatment testing data, demonstrating that Model 3 and, in particular, Model 4 are most consistent with the observed dynamics. As such, we concluded that PARPi response involves both drug-dependent and independent steps. Bars denote the r^2^ value of the prediction based on the maximum likelihood parameter estimates and error bars denote 95% confidence intervals obtained from parametric bootstrapping. **c)** Maximum likelihood parameter estimates (solid bars) and confidence intervals (error bars) for Models 2-4.

**Figure S4.**
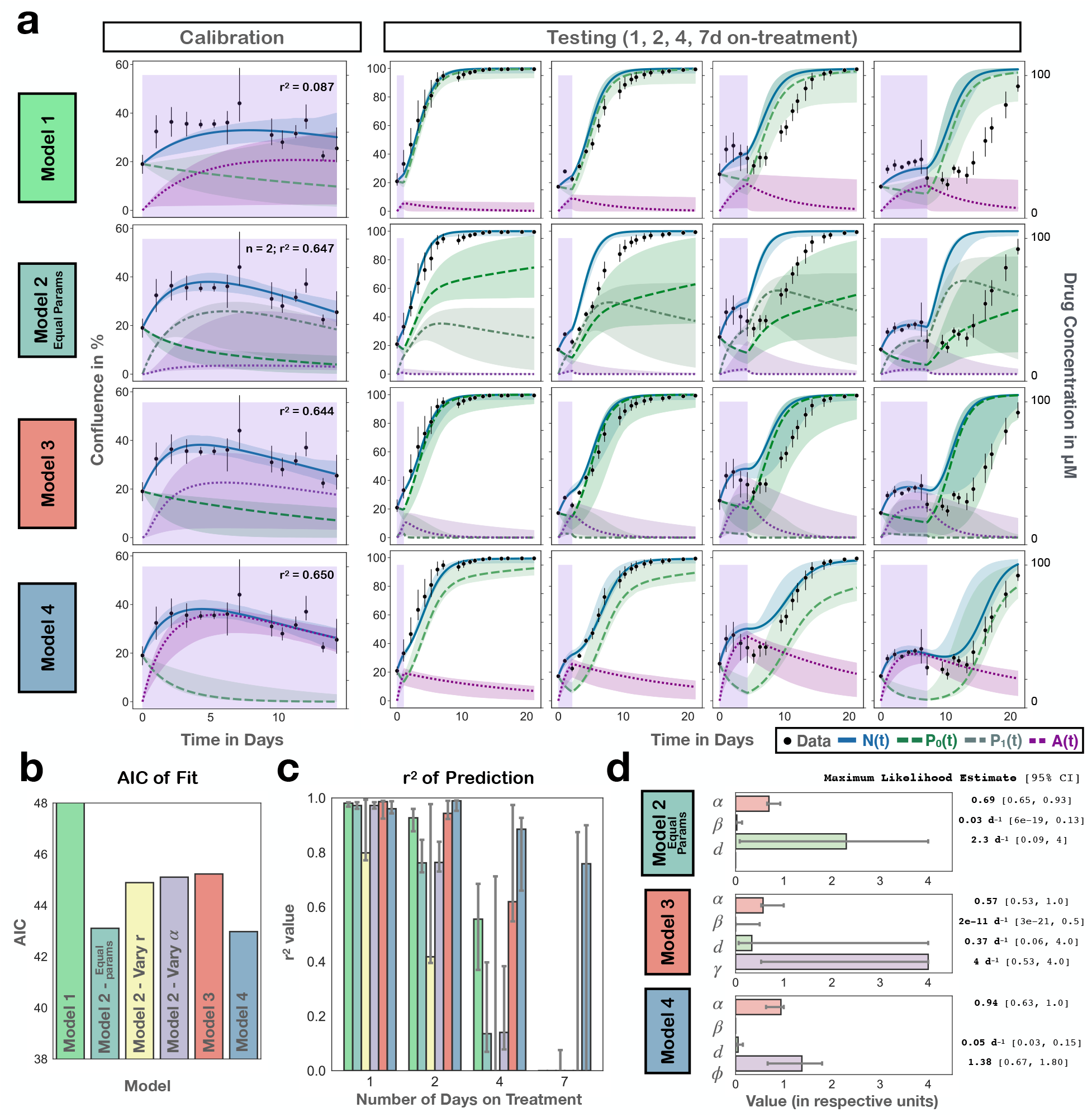
Repeating the treatment model analysis with the OVCAR4 cells confirmed that treatment response involved both drug-dependent and independent steps and corroborated the descriptive and predictive power of Model 4. **a)** Model fits (left column) and model predictions (right columns) for the four different models examined. For Model 2 we found that similar to the OVCAR3 data a value of n =2 yielded the best fits. Points and error bars denote mean and 95% CIs of observed confluence (3 independent replicates per condition). Lines depict the model predictions based on the maximum likelihood estimate, and bands indicate 95% CIs calculated via parametric bootstrapping. Note that due to issues caused by the existence of local optima in the likelihood surface, we turned off randomization of the initial parameter guesses when fitting Model 4. **b)** Quantification of the goodness-of-fit of each model. For reference, we also show the AIC of Model 2 when allowing r or alpha to vary as damage is accumulated. As for the OVCAR3 cells, this does not improve the fits. To allow for better comparison of Models 2-4, we cut off the y-axis at a value of 48. The AIC for Model 1 is 56.4. **c)** Quantification of the prediction accuracy of each model on the testing data. Bars denote the r^2^ value of the prediction based on the maximum likelihood parameter estimates and error bars denote 95% confidence intervals obtained from parametric bootstrapping. **d)** Maximum likelihood parameter estimates (solid bars) and confidence intervals (error bars) for Models 2-4.

**Figure S5.**
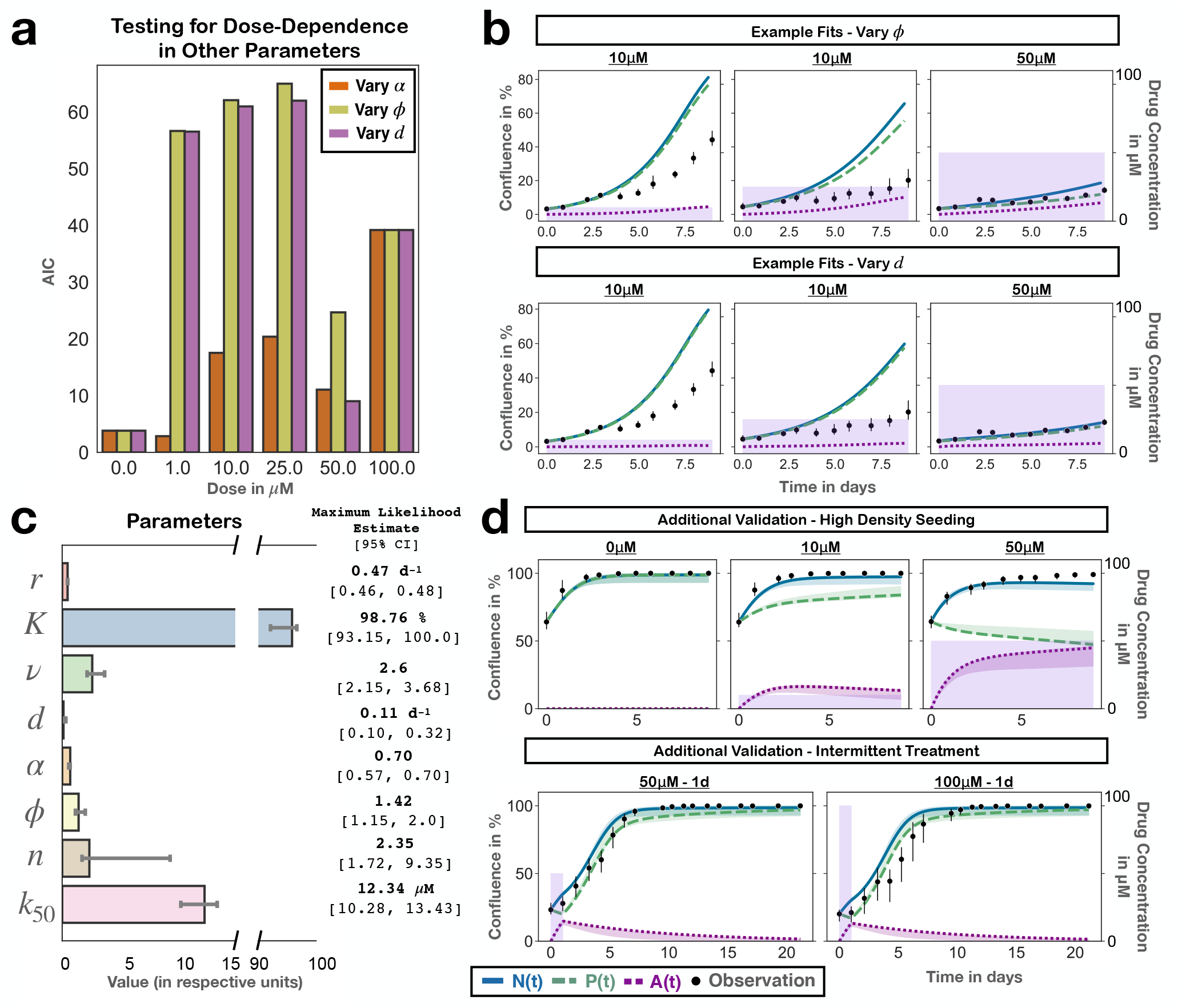
Additional details on the calibration and validation of the final model for OVCAR3 (Model 5). Throughout the figure, points and error bars denote mean and 95% CIs of observed confluence (3 independent replicates per condition). Lines or bars depict the model predictions based on the maximum likelihood estimate, and bands or error bars indicate 95% CIs calculated via parametric bootstrapping. **a)** Model 5 assumes that *α* varies with dose. To test the alternative hypothesis that one of the other treatment related parameters may vary with dose we fitted the model allowing either ϕ or d do be dose-specific instead (analogous to the protocol in Figure 5a). This plot compares the goodness-of-fit of the different models, showing clearly that a dose-specific *α* provides the most consistent explanation of the data (smaller AIC is better). **b)** Example fits from the analysis in illustrating how changing ϕ or d fails to produce a model that can describe the dynamics across different drug concentrations. Together, a) & b) corroborate our choice of a dose-specific *α* in Model 5. **c)** Parameter estimates for the final model. **d)** Model predictions for additional experimental conditions not shown in Figure 5 further corroborating the high predictive power of this model.

**Figure S6.**
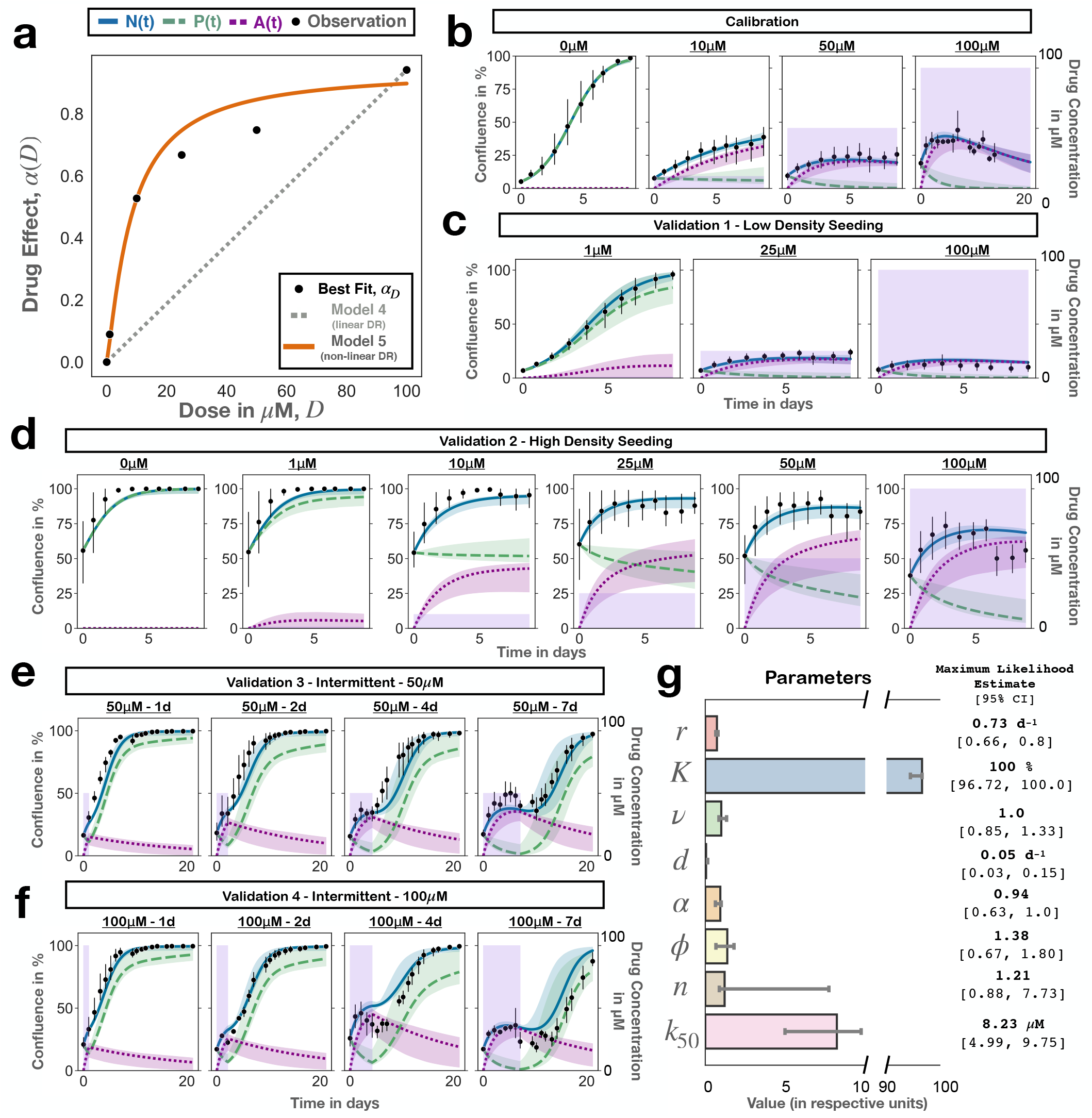
Calibration and validation of Model 5 for OVCAR4. Throughout the figure, points and error bars denote mean and 95% CIs of observed confluence (3 independent replicates per condition). Lines or bars depict the model predictions based on the maximum likelihood estimate, and bands or error bars indicate 95% CIs calculated via parametric bootstrapping. Note that because of issues with local maxima in the likelihood surface we did not randomize the initial parameter estimates when fitting to the bootstrap samples but instead started fitting from the Maximum Likelihood estimates. **a)** Empirical dose-response relationship derived from the data using the protocol from Figure 5a. Similar to OVCAR3, this shows a clear concave relationship, which cannot be described by the linear dose-response model assumed in Model 4 but is fitted well by the Hill equation assumed by Model 5. **b)** Fits of Model 5 to the calibration data for OVCAR4. **c)** Model predictions for the treatment dynamics under continuous treatment at different drug concentrations for cells seeded at a low density. **d)** Model predictions for continuous treatment at different drug concentrations for cells seeded at a high density. **e)** Model predictions for the response to increasing lengths of treatment at 50µM followed by treatment withdrawal. **f)** Model predictions for the response to increasing lengths of treatment at 100µM followed by treatment withdrawal. Together, Panels b)-f) demonstrate that Model 5 also provides highly accurate fits and predictions for the treatment response of OVCAR4 cells. **g)** Estimated model parameters.

**Figure S7.**
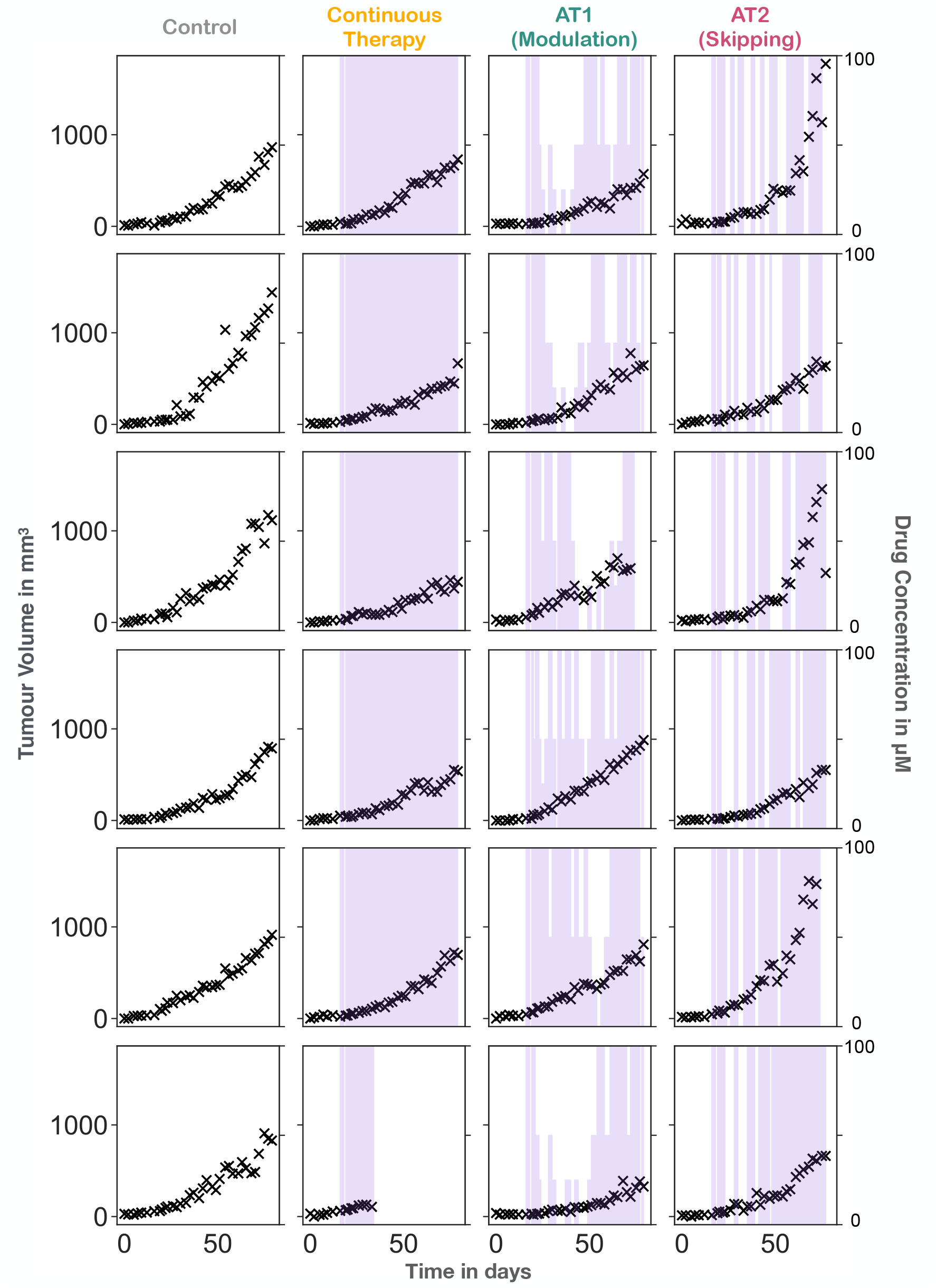
Treatment trajectories of each individual mouse during the in vivo experiment.

